# Cilia dynamics creates a dynamic barrier to penetration of the periciliary layer in human airway epithelia

**DOI:** 10.1101/2025.03.21.644548

**Authors:** Erika Causa, Debasish Das, Luigi Feriani, Jurij Kotar, Pietro Cicuta

**Affiliations:** Department of Physics, Cavendish Laboratory, University of Cambridge, J.J. Thomson Avenue, Cambridge CB3 0HE, UK; Department of Mathematics and Statistics, University of Strathclyde, Livingstone Tower, 26 Richmond Street, Glasgow G1 1XH, UK

**Keywords:** Airway cilia, Mucociliary Clearance, Periciliary Layer, Metachronal waves, Ciliary coordination

## Abstract

The ciliated epithelium of the human respiratory tract is covered by the airway surface liquid (ASL), a protective fluid consisting of two layers: the periciliary layer (PCL), where motile cilia reside and generate fluid flow, and an overlying mucus layer. The complex structure and stratified nature of the PCL complicate both the prediction and quantification of fluid flow at the scale of individual or small groups of cilia, making it difficult to connect microscopic flows to macroscopic clearance. To tackle this challenge, we developed a novel methodology that involves ‘un-caging’ a fluorescent compound to trace the flow field within the PCL. Fluorescence is activated at micrometric spots within the cilia layer, and displacement vectors and diffusion are recorded using high-speed video. Our experiments reveal a complex fluid transport pattern, with displacement velocity along the epithelial surface varying due to a non-uniform vertical flow field. Additionally, we observed that cilia expel fluid at their tips, a mechanism likely aimed at preventing pathogen access to the epithelium. Simulations, where cilia are modeled as arrays of rigid rods with length asymmetry, support these findings and offer new insights into the dynamics of fluid transport in the respiratory tract and the critical role of cilia coordination.

**Significance Statement:** This study introduces an experimental pipeline to investigate fluid velocity and diffusion within the PCL of the human respiratory tract. By integrating experimental data with simulations of ciliary motion, we offer a robust framework to understand how cilia, depending on their collective beating properties, propel periciliary fluid in this structurally and dynamically complex environment. Our findings significantly expand the understanding of ciliary function, revealing that when cilia are beating coherently near cilia tips fluid is actively driven away from the epithelial surface. This suggests that coordinated cilia movement not only plays a key role in maintaining respiratory health by clearing mucus, but may also provide a dynamic barrier against pathogen entry.

---

**F**luid flow generated by the coordinated motion of cilia in the human airway epithelium underpins mucociliary clearance (MCC), a vital physiological defense mechanism (1–4). The efficiency of MCC is thought to rely on the interplay of three key elements: motile cilia that beat in coordination to generate the primary driving force; a low-viscosity lubricating periciliary layer (PCL) that allows efficient ciliary movement; and a viscoelastic mucus layer that traps inhaled particles, preventing them from reaching the epithelial cells. According to the currently accepted model of MCC, the tips of the cilia penetrate the mucus layer during the power stroke but do not interact with it during the recovery stroke (5). In this picture, the cilia act on the lower surface of the mucus sheet, propelling it unidirectionally along the airway towards the mouth and digestive tract (6, 7). Failure in one or more components of this system can lead to or exacerbate severe infections and chronic inflammatory conditions, such as in Cystic Fibrosis (CF), Primary Ciliary Dyskinesia, or Asthma (8–10). Unraveling the mechanisms that clear inhaled particles and pathogens from the airways is essential for understanding respiratory health and disease. Several factors influence mucociliary transport in the respiratory system, including the density, spatial distribution, orientation, length, ciliary beat frequency (CBF), and stroke amplitude of the cilia, all of which affect ciliary coordination (11). The characteristics and rheological properties of the PCL and mucus also interact with these factors (12). Despite extensive advances in experimental and modeling techniques, our understanding of the fundamental mechanisms of airway mucociliary transport remains fragmented (13).

Mucus is a solution of mucin proteins, suspended in a matrix of charged ions and DNA fragments (14). Its composition is tightly regulated by the epithelium and can change dramatically in the context of inflammation or genetic diseases such as cystic fibrosis (CF) (15). The structure and composition of airway mucus have been extensively studied and are relatively well understood (16, 17). Standard methods, such as the use of tracer beads, allow precise tracking of mucus flow and clearance patterns at the macroscopic scale, both *in vitro* and *in vivo* (18–20). However, probing mucus at microscopic scales and characterizing its rheological properties *in situ*—including frequency-dependent viscosity and elasticity—remain significantly challenging (21–23).

The PCL has traditionally been described as a simple liquid-filled domain in which the cilia beat (24). However, this model failed to explain why the inter-ciliary space is impenetrable to nanometric objects such as beads or the macromolecular gel-forming mucins found in the mucus layer. In 2012, Button *et al*. (25) proposed an alternative hypothesis: the PCL acts as a ‘polymer brush,’ formed by membrane-spanning mucins grafted to the surface of the cilia, with a mesh size of 20-40 nm. This architecture stabilizes the PCL against compression by the mucus and serves as a barrier, restricting particle access to the cell surface (26). The intricate, impenetrable nature of this structure complicates the use of conventional tracer beads to quantify fluid flows within the PCL.

Matsui *et al*. overcame this limitation by labeling the airway surface liquid (ASL) of well-differentiated tracheo-bronchial cell cultures with caged fluorescein-dextran molecules (27). By measuring the displacement of the dye, which they photoactivated in columns perpendicular to the epithelium, they discovered that the PCL was transported at the same rate as the mucus (39.2±4.7) µm) when both layers were present. However, transport slowed down significantly to (4.8±0.6) µm upon mucus removal, suggesting that PCL movement was largely driven by friction with the overlying mucus layer. This method, however, only yielded the average PCL velocity, leaving the exact mechanisms of fluid transport within the PCL unclear.

Recent experiments using explanted tracheae from both mice (28) and humans (29), where the mucus was removed and replaced with a watery physiological solution, demonstrated effective transport of beads several hundred micrometers away from the epithelium. These findings confirmed that ciliary motion can drive fluid transport even in the absence of mucus. However, these studies did not specifically measure fluid flow within the ciliary layer itself.

Mathematical modeling of cilia has been fundamental in understanding the fluid flow generated by their beating. One of the earliest hydrodynamic models, developed by Blake (1971) (30), introduced the concept of a spherical envelope around closely spaced cilia, effectively coarse-graining the dynamics of individual cilia. While these continuum models are useful for studying large-scale phenomena, they do not resolve flow fields around individual cilia. Consideration of fluid flow at the level of individual cilia or small patches is important for optimized fluid transport and enhanced clearance, as evidenced by recent studies (31, 32). Numerical methods such as the immersed boundary method (33), multiparticle collision dynamics (34), and slender body theory (35, 36) have been developed to simulate individual cilia. In parallel, reduced-order models have been developed to explore synchronization and metachronal wave formation. For example, Niedermayer *et al*. (37) introduced a phase oscillator model where each cilium, represented by a spherical bead, follows a circular trajectory with a variable radius, allowing for hydrodynamically induced synchronization. This spherical bead model has been extensively used to address various questions on phase synchronization from a dynamical systems perspective (38–40). In this study, we do not focus on the internal molecular mechanisms driving synchronization or generating metachronal waves but instead on accurately computing the fluid flows generated by an array of cilia. Prescribing the kinematics of each cilium, we fix the wave pattern for the entire array; slender body theory can then resolve fluid flow around individual cilia while accounting for hydrodynamic interactions. Computing flow fields allow us to track the trajectories of passive particles introduced into the simulation domain, and compare to experimental observations.

In this paper, we perform at the same location high-speed videos of bronchial cilia beating in a Newtonian fluid and measurements of fluid transport in the PCL, using caged dyes. Caged dyes are small synthetic fluorophores covalently bonded to a photo-labile protecting group, which makes them non-fluorescent until exposed to specific wavelengths, typically UV (41, 42). This feature provides precise spatial and temporal control, making caged dyes ideal for tracking and mapping flow fields (43, 44) and cellular activities in targeted regions (45). The dye is activated in micrometric spots in the cilia-beating plane and the fluid speed is measured by tracking dye displacement along lines parallel to the epithelium. Quantitative velocity fields are resolved down to the micron scale by repeatedly activating the dye at different positions in the PCL and assessing the speed at each point. Our results reveal a complex fluid transport pattern: the displacement velocity along the epithelial surface varies due to fluctuations in the vertical component of the flow field. This variability underscores the intricate dynamics of ciliary-driven fluid movement. We compare these experimental results with simulations of ciliary arrays, providing a robust framework for understanding how fluid is propelled within this densely structured environment. We also observe a sub-diffusive spread of the activated dye in the ciliary layer. Significantly, these findings suggest that by actively propelling fluid vertically toward the airway lumen at their tips, cilia may help preventing pathogens from reaching the cell surfaces. This dual function highlights the critical role of ciliary movement in maintaining respiratory health and preventing disease.

## Results

### Visualization of Fluid Flow in the PCL using a photo-activated caged compound

The rigid plastic membrane supporting a well-differentiated human bronchial epithelium can be manipulated to image cilia from the side, so to see their beating pattern, as described in (46). The liquid overlaying the PCL then consists of a solution of 90% PBS and 10% culture medium with caged-fluorescein dye 1-2 mM. See *Methods* section for full details. For each experiment, a field of view (FOV) containing a few multiciliated cells was recorded using bright field microscopy for 5 seconds, at a rate of 80 frames per second (fps). This allowed us to calculate the cilia beating frequency (CBF) and to delineate the epithelium contour. We chose to image only cells whose beating patterns were oriented within the imaging plane, to the degree that this is clear by eye from how cilia remain in focus. Following the initial recording, the fluorescein dye was activated using a 12 ms pulse of 405 nm collimated light, focused through a 60× water immersion objective (NA = 1.2, Nikon) on a spot within the PCL with radius *<* 1.5 µm (Figure 1.A-C). The FOV was subsequently recorded in 5-second epifluorescence microscopy videos at 120-150 fps. The recording was started 0.30 s prior to the dye’s activation. After photo-activation, the dye is displaced and spreads out as a consequence of cilia-driven flow and diffusion (Figure 1.D-E). We measured the center of mass (*d*) and the width of the fluorescent patch (*σ*) as it moved along a measuring line parallel to the epithelium (Figure 1.F) and segments perpendicular to the epithelium.

**Fig. 1.**
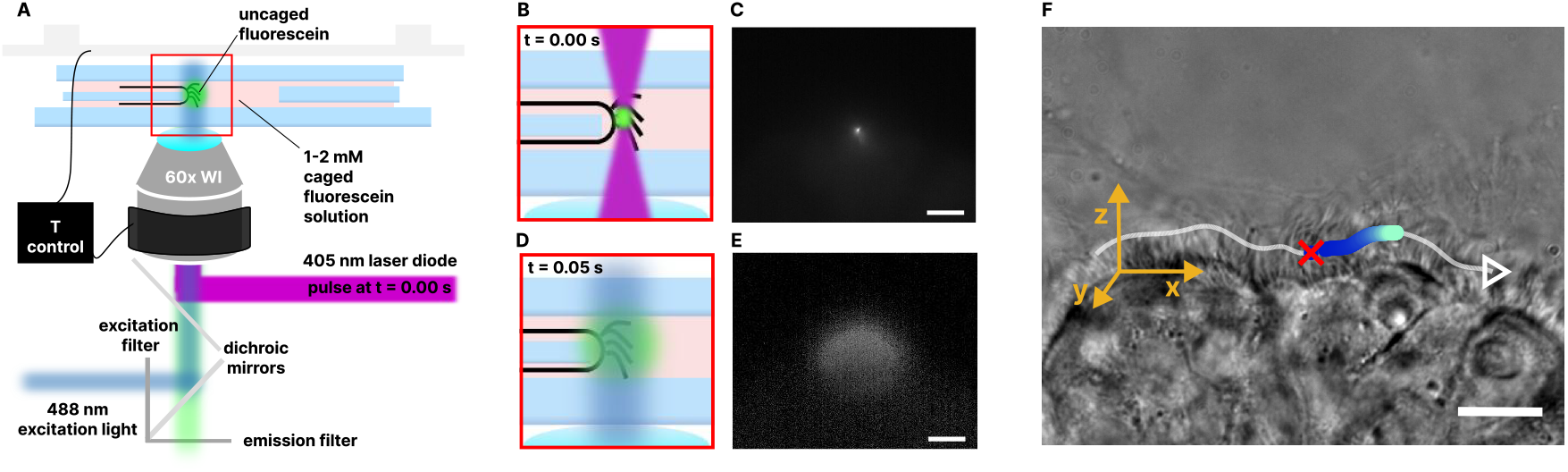
Experimental pipeline to probe PCL flows. (A) Schematic representation of the optical and sample set-up. The flow field in the PCL is probed by releasing a fluorescent dye from a caged state, activating it with a 12 ms pulse of collimated 405 nm light from a laser diode, focused through the objective. (B,C) The dye is released in a circular spot with radius *<*1.5 µm. (D,E) Immediately post-pulse, the fluorescent dye diffuses and is transported by the beating of cilia. (F) The dye displacement (time, blue to light blue) is measured along a measuring strip (indicated by the white line). The strip is at constant distance from the epithelium, orientated in the cilia’s power stroke direction, and passes through the dye activation spot (red cross). Scale bars are 10 µm.

### Velocities Induced Across PCL by Ciliary Beat

For each experiment, the fluid transport is measured as a function of the dye activation position within the PCL from the base to the apex of cilia. This is achieved by photo-activating the dye in different locations, chosen distant at 1.5 µm intervals from each other, progressing from the cell apical surface to the ciliary tips. Cilia are typically 7-9 µm long (2), so 5-7 positions are usually recorded. For each dye activation site, either a simple linear regression or a segmented linear regression is fitted to the displacement data. In the simple linear scenario, a single constant speed is derived from the slope of the fitted line (*SI Appendix*, Fig. S7). In the segmented approach, a short-time and a long-time speed are determined from the slopes of the segments before and after a break-point, which is set by eye (figure 2.A). This break-point indicates the midpoint of the transition between shortand long-time regimes, and is typically observed around 0.4 s, though sometimes up to 1 s after activation. This varies across FOVs. The dual-speed transport is detected in 87% of the experiments and also observed in simulations. This phenomenon can be attributed to changes in the vertical component in the flow field, as evidenced in both experimental (Figure 2.C) and simulated data (Figure 5).

**Fig. 2.**
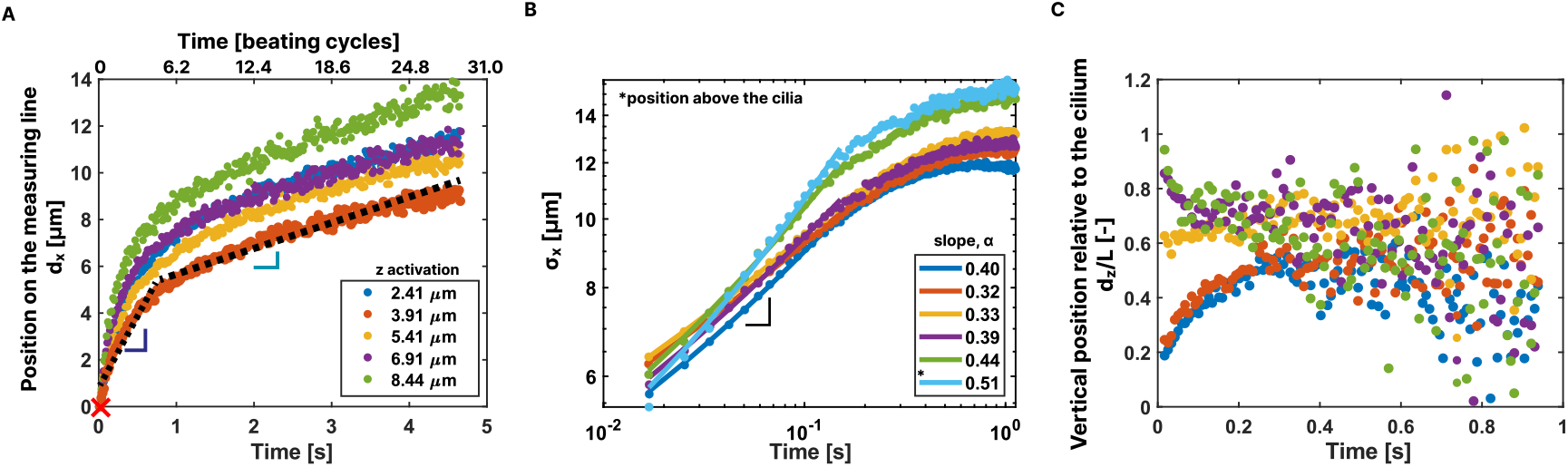
Advection and diffusion of the dye are measured. (A) The displacement along lines parallel to the epithelium reveals two distinct transport speeds in most experiments, termed as ‘short-time’ (blue gradient) and ‘long-time’ (light blue gradient) regimes. The transition is not sharp, but typically lies between 0.4 and 1 s (displacement and time are relative to the location and moment of activation). The short-time velocity is always the larger one, as explained in the text. The dye displacement is recorded in 5 s videos; in the case of the experiment in this figure, where the average measured *CBF* was (6.2 ± 1.4) Hz, this corresponds to 31 beating cycles (see upper axis). (B) As well as being advected, the dye spreads along the measuring line, quantified as the standard deviation of the Gaussian fit of the fluorescence intensity profile over time (*σx*). The logarithmic scale plot of this spread reveals sub-diffusive behavior across different distances from the epithelium. This sub-diffusivity is more pronounced at the cilia base as the power law exponent *α* increases moving towards the liquid bulk. Normal diffusivity (*α* = 0.5) is observed above the cilia layer (last position) - see *Methods* for details about diffusion fits. (C) For each frame, the dye displacement is measured also along *z*, as the position of the mean of the Gaussian that fits the fluorescent profile, along vertical lines positioned such that their *x* coordinate corresponds to the center of the dye cloud at that time, measured on the measuring line, *dx*(*t*). Generally, when activated within the ciliary layer, the dye is pushed towards the cilia tips. Variation in the vertical component of the flow field causes changes in the transport rate along the epithelium.

In Figure 3.B, the measured speeds are averaged and plotted as a function of the dye activation position along the cilium, as illustrated in Figure 3.A. The dye activation height along the cilium, *z*, and the velocity, *U*, are non-dimensionalized with respect to the cilium length, *L* (µm), and the ciliary oscillatory velocity, *CBF* × *L* (µm*/*s), respectively. This normalization enables the aggregation of multiple experiments into a unified dataset. For each region imaged, the cilium length is measured as the distance between the apical surface of the epithelium and the cilium tips at the *x*-coordinate where the dye is activated. The *CBF* is determined for each FOV from bright-field videos using an algorithm based on Fast Fourier Transform (FFT), as described in the *Methods* section. To account for potential variability in cilium length and beating frequency across different cells and experimental conditions, normalization is performed separately for each FOV. Cilia average length and *CBF* across all the experiments are (8.4±0.3) µm*/*s and (5.6±0.4) Hz, respectively.

**Fig. 3.**
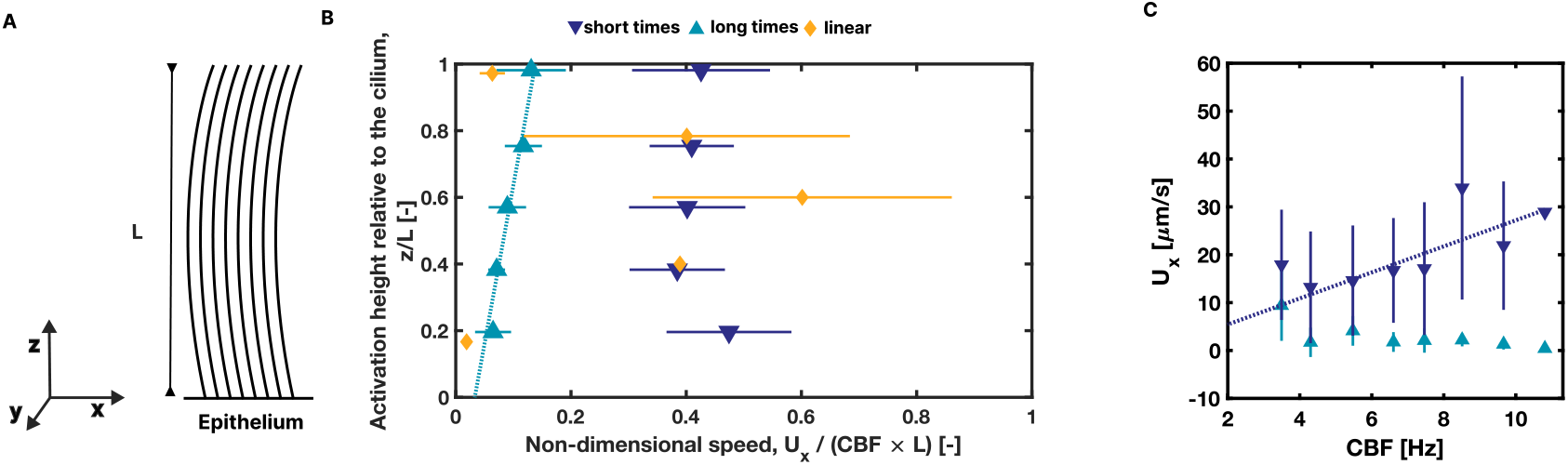
Short- and long-time velocities in the PCL. (A) Sketch of cilia from a ‘side’ view, the (x,0,z) plane in this work. (B) Average fluid velocity profiles in the PCL at 37 °C. Short-time fluid velocity (▾) remains relatively constant across the PCL, while long-time velocity (▴) increases with increasing distance from the epithelium. The increasing trend is captured well by a weighted linear least-squares fit (dashed light blue line). These velocity profiles represent averages obtained from 91 fields of view and 3 inserts across 3 experimental sessions. The measured average long-time velocity is (3.79 ± 1.12) µm*/*s, which is in close agreement with the findings of Matsui *et al*., who reported similar velocities under mucus-free conditions (27). In about 13% of the experiments, a linear displacement plot is observed. The corresponding linear speeds, represented in yellow (♦), have the same order of magnitude as both the short-time and long-time speeds but do not specifically belong to either case. (C) Short-time and long-time speeds measured across all experiments are binned into eight groups and plotted against *CBF*. The short-time speed data points are fitted with a weighted linear fit (dark blue dashed line) constrained to pass through the origin, showing a strong correlation with *CBF*. This suggests that immediate ciliary activity plays a key role in determining short-time speeds. In contrast, long-time speeds exhibit a more complex dependence on *CBF*, indicating that additional factors influence transport dynamics over extended observation periods.

Across the cilia layer, the non-dimensional short-time speeds were found to maintain consistent values. The long-time speed changes with distance from the epithelium: Figure 3.B shows the data is well fitted linearly (free intercept), R^2^ = 0.96. Notably, the short-time speed exceeds the long-time speed by a factor of 5.5±1.8. The minority of conditions with a single speed do not seem to belong specifically to either cases, and their speeds take values in the range of the short and long-time. Speed averages are (18.8±3.4) µm*/*s for short-time, (3.8±1.1) µm*/*s for long-time, and (9.9 ± 3.0) µm*/*s for the cases with a constant velocity.

### Correlation between PCL speed and CBF

The efficiency of the airway clearance mechanism is intrinsically dependent on the speed at which cilia beat, which varies significantly depending on several factors, including the health status of the individual, the location within the respiratory tract, and external conditions during experimental assessments. CBF values have been reported from about 3.1 Hz to 19.7 Hz in human tracheo-bronchial cells, influenced by physiological factors like pH, temperature, and humidity (47). Ideal conditions for MCC have been identified at temperature of 37 °C with 100% humidity, under which the PCL maintains its physiological height, thereby optimizing CBF and MCC (48). At lower temperatures and humidity levels, the PCL height decreases, reducing CBF and slowing MCC (2). Additionally, CBF can be modulated by drugs and environmental stimuli such as ATP, which enhances CBF by increasing intracellular calcium levels (49). Given this, determining how variations in CBF influence the transport dynamics of the PCL is essential for understanding MCC.

To investigate the relationship between the speed measured along the *x*-axis and the recorded *CBF*, we binned short-time and long-time speed data into eight groups and plotted them against *CBF*, observing an increasing trend of short-time speed with *CBF*. To quantify this dependency, the short-time speed data were fitted with a weighted linear fit, visually represented by a blue dashed line in Figure 3.C, yielding a slope of 2.72 µm. These findings highlight a strong correlation between short-time speeds and *CBF*, indicating that these velocities are likely influenced by immediate ciliary dynamics. In contrast, long-time speeds show a more complex relationship with *CBF*, suggesting that other factors contribute to transport dynamics over longer observation periods. We explore this relationship further in *SI Appendix*, section 5.

### Oscillatory fluid displacement

In a minority of instances, time-oscillatory patterns have been observed in the dye displacement plots for short-times after the dye activation (up to 1.2 s). These oscillations in PCL fluid transport are likely to reflect the observations of (50), and to be indicative of particular coherence in the phase-beating properties of cilia at the point of the dye release. The detection of these oscillations predominantly at short timescales is attributed to the small size of the dye patch during these intervals, which allows for the signal to probe a parcel of fluid with spatially coherent dynamics. In an effort to quantify these observations, the regions where oscillations were evident, denoted by the zoom in Figure 4.A, are fitted to determine the amplitude and period of the oscillatory displacement (additional info provided in *SI Appendix*, section 6). For each of these experiments the oscillation period is calculated at different distances from the epithelium (coloured squares in Figure 4.B), and can be compared to the *CBF* obtained via FFT analysis, depicted by black squares. The correspondence between these two measurements was found to be remarkably consistent over seven experiments, reinforcing the hypothesized link between ciliary activity and the observed fluid oscillations.

**Fig. 4.**
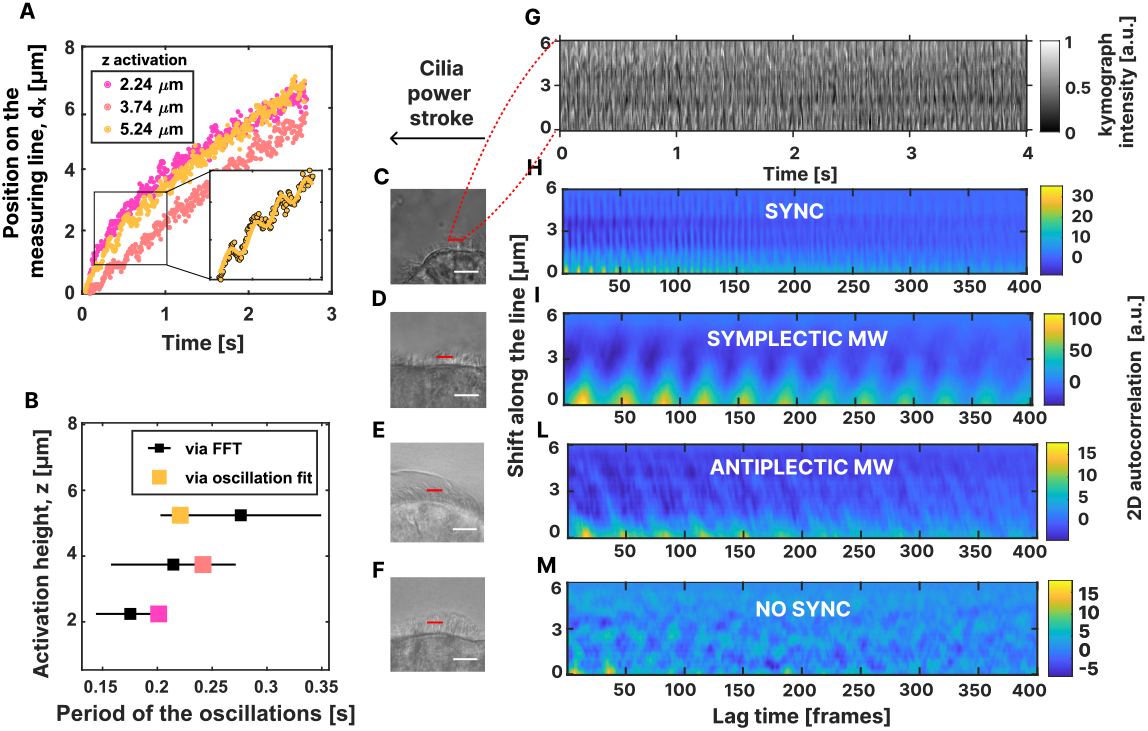
Temporal Analysis of Oscillations in cilia and Dye Displacement. (A) Emergence of oscillatory fluid flow at short timescales, observed at three different dye activation points from the epithelial surface. In the specific time interval where the dye displacement exhibits oscillatory behavior, a sinusoidal curve fitting is applied to the data to measure amplitude and period of the oscillations (*SI Appendix*, Supplementary Fig.10). A zoom highlights the fit (yellow curve) and the corresponding data points (dots). (B) The oscillation periods extracted from the fits are plotted along with *CBF* values determined through FFT analysis, denoted by black squares. The congruence between the measured periods and *CBF* data validates the hypothesized connection between the ciliary motion and the observed fluid oscillations. (C-F)Visualization of 100 pixel long segments employed for ciliary correlation analysis in the PCL. The segments are oriented in the direction of the power stroke. (G) Grey-scale intensity kymographs are generated on these segments in the videos, choosing examples with and without evident oscillatory fluid flow. (H-L) 2D autocorrelation plots of the kymographs show correlation between oscillatory fluid flow and parallel lines of spatio-temporal coordination. (M)The high-correlation parallel lines are absent when the fluid displacement is not oscillatory. Video recording frame rate was 120 fps in the synchronous case and 150 fps in the other three cases. Videos illustrating key exaples of the four synchronization scenarios are available in the *SI Appendix*, Supplementary Movie S1.

The emergence of oscillations in the PCL fluid flow and their relation to ciliary motion are also investigated using greyscale intensity kymographs along specific segments within the ciliary layer, as detailed in the *Methods* section. By employing cross-correlation analysis of these intensity measurements, we could evaluate the spatio-temporal coordination of ciliary activity. Our comparative analysis focused on videos that depicted oscillatory dynamics in the dye versus those that did not and revealed that the presence of oscillations is marked by parallel lines of high correlation in the 2D autocorrelation plots in Figure 4.H-L. These lines indicate a coordination in ciliary beating: vertical lines suggest cilia beating in unison (in-phase beating) while angled lines are indicative of a consistent phase difference between the beats of adjacent cilia, forming a metachronal wave (MW) pattern, as shown in (51). Conversely, the absence of such high-correlation lines in the autocorrelation plots (see Figure 4.M) points to a lack of coordinated beating—cilia beating asynchronously, each in a different phase. This behaviour was consistently observed in FOV where the dye did not exhibit oscillatory transport.

### Metachronal waves speed and wavelength

Adjacent cilia in dense arrays usually do not beat in phase, but maintain a constant, small phase difference, resulting in a propagating pattern called metachronal waves visible at the surface formed by the cilia tips. Symplectic MWs move in the same direction of the power stroke and occur when the phase lag between two adjacent cilia is negative. Conversely, a positive phase difference results in antiplectic MWs that travel opposite to the flow direction. Antiplectic waves have been shown to enhance fluid transport in ciliary carpets (52, 53).

In our study, we focused on two experiments where the phase lag between neighboring cilia was detected to measure the MW speed. We examined the behavior of auto-correlation against lag time at a constant space shift (*SI Appendix*, section 7.A). The data were fitted using a damped oscillator equation. The speed of MW propagation measured was (0.79±0.18) mm*/*s for the symplectic wave (Figure 4.I) and (0.17±0.08) mm*/*s for the antiplectic wave (Figure 4.L). These speeds align with previously measured values in mammalian trachea, typically ranging between 0.1 and 0.4 mm/s (54).

To measure the MW wavelength, we extended our analysis to the maximum segment length within the cilia patch and examined the behaviour of autocorrelation against space shift at specific lag times (*SI Appendix*, section 7.B). We determined the wavelength *λ* as the distance between two consecutive peaks (or valleys) in the autocorrelation vs space shift plots. The measured wavelength averaged (13.4±0.8) µm across the two experiments; This value is comparable to a coherence scale previously measured in our group (*λ* ≈ 20 *µm*) in 2D cilia carpets (55) and also to the coordination scale one expects from a balance of hydrodynamic coupling to thermal noise (56).

### Simulation results

The fluid dynamics within the PCL of the human respiratory tract are governed by Stokes flow due to the low Reynolds number in this microenvironment, where viscous forces dominate and inertial effects are negligible (30, 57–60). To simulate the cilia-driven fluid flow, we employ slender body theory (SBT), a mathematical framework ideal for modeling elongated structures like cilia (35, 61–65). SBT captures the hydrodynamic interactions between different parts of the cilium as well as the interactions between multiple cilia. SBT is essentially an integral equation relating the velocity of the slender body to the entire force per unit length distribution over the body. An important consideration in our simulations is the rigid epithelial surface to which the cilia are attached which imposes a no-slip boundary condition on the fluid. To model this, we used a modified Green’s function in the integral equation that inherently satisfies the no-slip condition at the epithelial wall, obviating the need to discretize the wall and thus simplifying the computational domain. This approach allowed us to accurately compute the hydrodynamic forces acting along each cilium by prescribing the kinematics. The wall is situated at *z* = 0.

The distance between each cilium base is denoted by *d*. The angle, *θ*, which each cilium forms with respect to the wall, is measured in the counter-clockwise direction. The cilia themselves are modeled as rigid rods that oscillate around the mean position of *θ* = *π/*2. The length of the cilia *L* = 10 µm is chosen as the primary length scale in the model. The length of the cilium is assigned to be *L*\* = 1 and *L*\* = 0.6 in the power and recovery stroke, respectively (see 5.A). This difference in length between the strokes introduces the necessary asymmetry in the beating pattern, which effectively pumps fluid in the positive *x*− direction. The numerical method developed here is an extension of our previous simulations where the cilia were static (66). Further details are provided in *SI Appendix*, section 8.

We first examined the trajectories of particles released at incremental heights of 0.1 from the wall, within the beating plane of a single cilium. Recognizing that the initial position of the cilium within its beating cycle will influence the results, so to match a mean flow we conducted multiple simulations releasing particles at various cilium positions in the cycle. We then averaged the trajectories of particles released from identical locations over one complete cycle. This method ensures that our results are robust and representative of the average particle behavior (see *SI Appendix*, section 8 for details). Remarkably, even a single oscillating cilium is able to reproduce qualitatively the key features of particle trajectories observed in experiments: the characteristic double slope along *x*, as most particles either get pushed away from the epithelial wall or get sucked towards it (see Fig. 5.B left). This phenomenon occurs because particle velocities sharply decrease as they approach or move away from the wall, leading to a reduction in their speed (see Fig. 5.B middle). The particles that do not change their height from the wall suddenly exhibit a much gentler slope in their trajectories along *x*. The trajectories of the particles in the *x* − *z* plane are plotted in (Fig. 5.B right). Some of the trajectories are characterized by oscillatory motions that closely mimic the experimental data shown in Fig. 4.A. In simulations, the trajectories are plotted as moving averages with a period of one complete oscillation of the cilium.

**Fig. 5.**
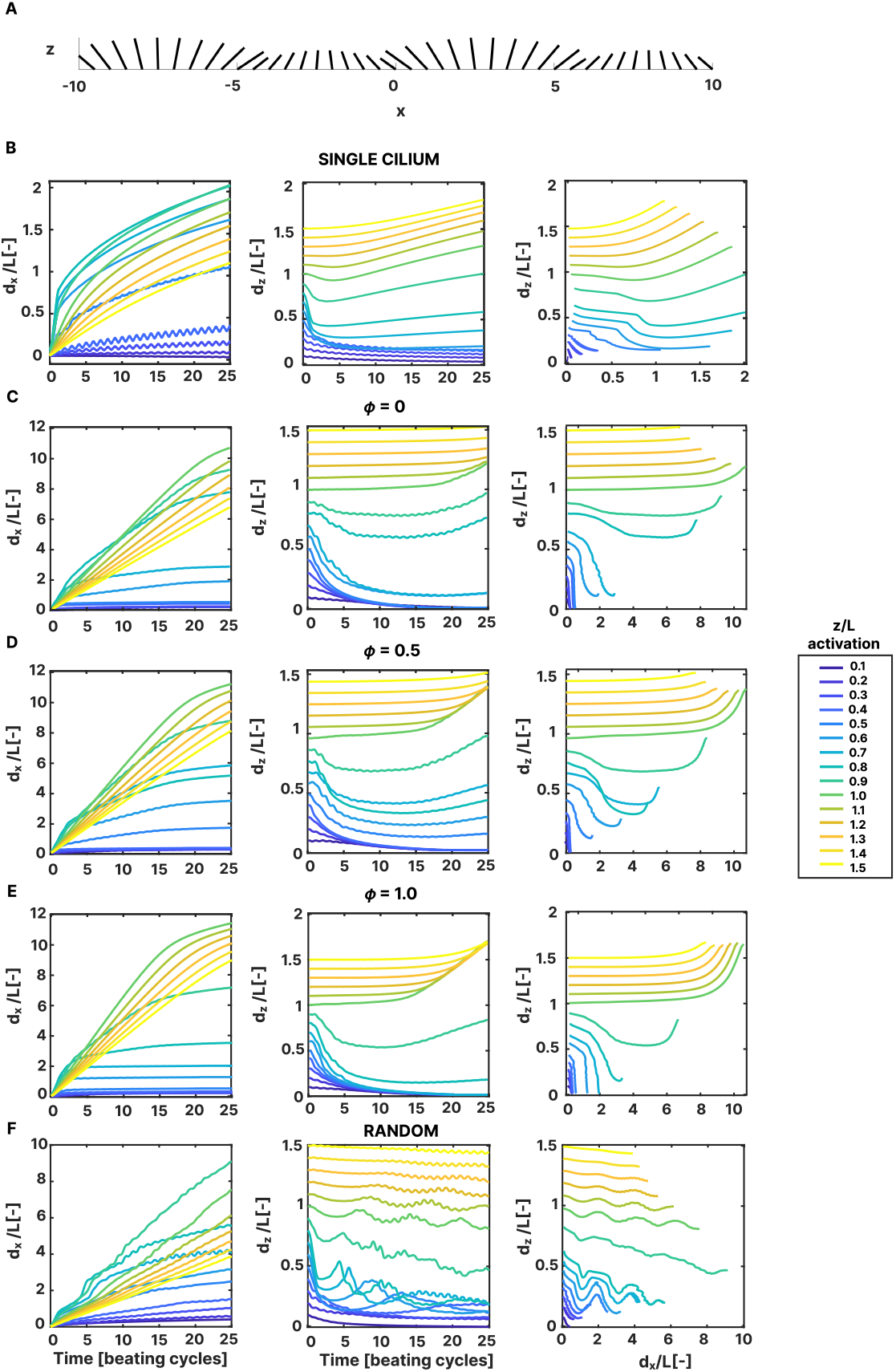
Simulations allow us to probe mechanisms of PCL transport generated by single cilia and ciliary arrays. (A) Schematic diagram of the numerical model: an array of cilia beating in a metachronal wave pattern; the phase difference between neighboring cilia is explored. (B-F) Displacements plots of particles released at incremental heights from *z* = 0.1 to *z* = 1.5 (*z* = *L* = 1 correspond to the extended cilium length). The simulated cilia transport fluid along the epithelial surface (*x*, left) and also perpendicularly to the epithelium (*z*, middle). Trajectories in the beating plane *x* − *z* (right) recapitulate one of the key experimental findings. Results are from (B) single cilium simulations, (C) ciliary array beating in synchrony (*ϕ* = 0), (D-E) cilia beating in a MW pattern: *ϕ* = 0.5 corresponds to a single MW propagating across the array, and *ϕ* = 1 corresponds to two MWs in the array; (F) cilia beating in a random pattern. In the simulations, the time interval explored has been chosen to be similar to the one of experiments – cilia beat at ~ 5 Hz (the average measured *CBF* at 37 °C is (5.6±0.4) Hz) and the video captured are ~ 5 s long so we show here cilia beating for ~ 25 cycles.

We then extended our simulation to a one-dimensional array of 41 oscillating cilia, exploring synchronization scenarios: beating synchronously, or with a constant phase difference between them, or with random phases differences between them (see Figure 5.(C-F)). The phase difference between neighboring cilia, *ϕ*, determines the synchronization pattern. When *ϕ* = 0, the ciliary array beats in perfect synchrony. For *ϕ* = 0.5 and *ϕ* = 1, the cilia exhibit a MW pattern, corresponding to one and two MWs propagating through the array, respectively. A video showing streamlines around the ciliary array with two MWs can be found in the *SI Appendix*, Supplementary Movie S2. All the particles are released in the cilia beating plane, *y*0 = 0. The dynamics observed in the single cilium simulations, characterized by wall suction and repulsion, are also found in the array simulation when the cilia beat synchronously (Figure 5.C) or metachronously (figure 5.D-E). These effects become more pronounced as we increased the phase difference between adjacent cilia, simulating two full metachronal waves (see Figure 5.E left). Particles released below *z* ~ 0.8 undergo suction, resulting in decreased velocities along the *x* axis, while those above this height are expelled, experiencing a subsequent velocity decrease. Remarkably, the distinctive double slope pattern along *x* observed in experimental data is consistently replicated in all simulation scenarios. Notably, the slope change due to wall suction occurs earlier, closely mirroring experimental observations, whereas the change due to expulsion is significantly delayed.

The vertical position of particles does not necessarily decrease or increase monotonically. This behaviour results from the finite size of the cilia array in our simulations. Additionally, particles may move out of the plane, further contributing to non-monotonic displacement. If simulations were conducted with an infinitely long cilia array in the x-direction and an infinite number of such arrays in the y-direction, we would expect a strictly monotonic displacement of all particles in the vertical direction. The velocity streamlines that gives rise to the complex trajectories in Figure 5.D are provided in supplementary material.

When particles are released in a plane parallel to the *x* − *z* plane, i.e. for a finite but small *y*, they are mostly repelled from the *x* − *z* plane depending on the height where they are released (see *SI Appendix*, section 8 for more details). In this case, particles released above 80% of the cilium length are still expelled, but the suction effect is dramatically reduced. Hence, the dual slope only exists due to the expulsion effect and occurs much later than that observed in experiments. This confirms that the sharp change in the velocity seen in experiment is due to a sudden change in the height of the particles from the epithelial surface, in the beating plane. We then assigned random phases to each cilium in the array, and particle trajectories were computed under these conditions in Fig. 5.F. Particles drawn towards the wall exhibited a change in slope along the *x* − direction as their velocities decrease. Interestingly, unlike the previous scenarios, no repulsion effect was observed in this configuration; particles released above *z* ~ 0.8 appear to get also sucked towards the wall (See Fig. 5.F middle). This is an undesirable characteristic from a biological viewpoint of the lungs, as foreign particles must be expelled from the PCL.

## Discussion

This study introduces an advanced experimental method for visualizing fluid flow within the PCL, allowing for precise mea-surement of cilia-driven transport at micron-scale resolution. By activating caged fluorescein in micrometric spots within the cilia’s beating plane, our technique significantly improves upon previous approaches, such as that of Matsui *et al*. (27), where dye activation occurred in broader 400 µm columns across the entire PCL height. This refinement enables a more detailed quantitative analysis of velocity fields, capturing fluid dynamics at precise locations within the PCL.

A distinct dual-speed transport pattern of the dye along the epithelial surface was observed in the experiments, consistent across different activation positions of the dye from the cilia base. The fluid speed measured shortly after dye activation (typically within 0.4 seconds) was approximately five times higher than the speed detected at longer intervals post-activation. This rapid change in the observed speed parallel to the epithelium can be attributed to complex average streamlines that are not strictly parallel to the epithelium, causing fluid parcels to acquire a vertical velocity component. The measured average long-time velocity was (3.8±1.1) µm*/*s which aligns closely with the findings of Matsui *et al*., who reported similar velocities in mucus-free scenarios ((4.8±0.6) µm*/*s) (27). The similarity between our average long-time speed and their measurements is expected, as their setup lacked the micrometric resolution along the *z*- axis provided by our method during the initial post-activation phase. Our approach offers enhanced visualization of local fluid dynamics, revealing a rapid initial transport of fluid parcels before a broader distribution along the *z*-axis. This means the long-time velocity is the net flow in the PCL; we are able to show a linear profile in z, which is consistent with a non-slip boundary approximately at the tissue plane.

We subsequently explored the impact of beating frequency on PCL transport by examining the correlation between the two measured speeds and the locally measured *CBF*. We found a significant dependency of short-time speeds on the *CBF*, suggesting that immediate ciliary dynamics play a key role in determining these velocities. In contrast, long-time speeds exhibit a more complex relationship with *CBF*, indicating that additional factors influence transport dynamics over extended periods.

To further investigate, we conducted some experiments at room temperature instead of 37 °C, and this led to a 20% reduction in *CBF*. The lower temperature caused a ~40% decrease in short-time transport speed and ~20% in long-time speed, indicating greater sensitivity of short-time speeds to *CBF* changes. These preliminary findings are discussed in *SI Appendix*, section 4.

Moreover, we investigated ciliary synchronization patterns by evaluating the spatio-temporal coordination of ciliary activity along segments within the ciliary layer. In experiments where time-oscillatory patterns were observed in the dye displacement plots, we identified two distinct beating patterns. The first, an in-phase beating pattern, was characterized by vertical lines of high correlation in the autocorrelation plot. The second, a MW pattern, was indicated by angled high-correlation lines. From these plots, we extracted MW speed wavelength, the second averaging (13.4±0.8) µm over two experiments. This value is approximately the width of a ciliated cell (~ 10 µm (67)) and similar to values of coordination length-scale previously measured in our group with multiscale-DDM (*λ*_*coord*_ ~ 20 µm) (55).

Further analysis of the vertical flow component revealed that in 84.5% of video recordings, cilia actively pushed fluid toward their tips, whereas in the remaining cases the fluid remained confined within the PCL. Specifically, the dye maintained the same *z* position in 13.4% of cases and was pulled toward the epithelium in only 2.1% of cases. These percentages were calculated by counting at each activation position whether *dz* was increasing, remaining constant, or decreasing, and then expressing each count as a percentage of the total number of activation positions across the entire dataset. Simulations of specific synchronization patterns— such as in-phase or MW beating—demonstrate an expulsion behavior where particles released above approximately 80% of the ciliary length are pushed toward the airway lumen. Foreign particles above this threshold would be expelled from the PCL, while those that penetrate below it might reach the epithelium. In contrast, simulations incorporating asynchronous beating do not exhibit this expulsion effect, increasing the likelihood that particles will be drawn toward the epithelium and potentially heightening the risk of infection. Although this correlation between ciliary synchronization and particle expulsion is clear in the simulations, it is not observed in our experimental data. Nonetheless, these findings underscore the crucial role of synchronized ciliary beating in protecting against pathogens.

Several aspects of our techniques warrant discussion. The complexity of sample preparation, particularly the removal of mucus, often poses significant challenges. Occasionally, cell detachment during the folding of the supporting membrane complicates imaging, resulting in unusable samples that undermine the reliability of outcomes. Additionally, even in well-prepared samples, identifying areas with densely packed, uniformly oriented cilia is difficult. This often leads to a scarcity of viable data points, requiring a greater number of experiments to achieve statistically valid results. Additionally, the use of caged dye presents challenges due to its rapid diffusion, limiting the duration for which the dye can be effectively tracked. Although dye diffusion is rapid, we confirm that mass transport dominates in our experiments, as the Peclet number is *Pe* = *UL/D* ≃ 1, with a characteristic length scale of around 10 µm, dye speed of 10 µm*/*s, and the diffusion coefficient of fluorescein in the PCL layer being *<* 10 × 10^−6^ cm^2^*/*s– that is the diffusion coefficient of fluorescein in a watery liquid, measured as described in *Methods*. Based on these considerations, the motion of larger objects like viruses would be dominated by advection (*Pe >* 1). To extend observation times, conjugating the caged dye with high molecular weight molecules like Dextran could slow the diffusion rate, but risks rendering the molecules too large to penetrate the nanometric mucin brush of the PCL. Our experimental setup excludes mucus to prevent visual interference, ensuring clearer imaging. However, this deviation from natural airway conditions limits the replication of real-world ciliary dynamics. Developing a method that allows precise PCL transport measurements in the presence of mucus would represent a breakthrough, offering closer approximations to *in vivo* conditions and potentially revealing new insights into mucociliary clearance (MCC).

Findings from this work suggest that cilia not only transport mucus but also act as a barrier, preventing viruses from reaching cell surfaces and actively removing pathogens from the ciliary layer. The protective role of active ciliary motion against viral infections has been observed, such as in the study by (68), where ciliostasis, or inhibited ciliary motion, increased susceptibility to H3N2 influenza infection. This dual functionality of cilia-driven flow in the PCL may explain the vulnerability of respiratory tissues to SARS-CoV-2, a virus that targets multiciliated cells and causes significant cellular alterations. Infected cells often exhibit dysfunctional, shortened, or misshapen cilia, as well as viral accumulation along membrane ruffles (69). These disruptions impair mucus transport and facilitate viral access to deeper tissues by negating the protective expulsion effect of motile cilia, exacerbating the infection. Our findings thus contribute not only to understanding PCL dynamics but also to broader implications of ciliary function and its disruption in respiratory health and disease.

## Materials and Methods

*In vitro* reconstituted Human Bronchial Epithelia MucilAirTM were purchased from Epithelix S`arl and cultured following the company’s recommended protocol. Velocity measurements were conducted in 15 experiments using 3 distinct inserts at 37 °C. All cells utilized for speed and diffusion measurement originated from a single healthy donor. Data for oscillation plots were sourced from 7 experiments, utilizing cells from two distinct healthy donors. Autocorrelation analysis was conducted on data from 8 experiments, five of which were selected from the oscillation data based on cilia patch shape and imaging clarity. These included three cases with no synchronization, one with full synchronization, three exhibiting symplectic MW, and one showing an antiplectic MW.

### Experimental set-up

Prior to experiments, a mucus wash was performed by incubating the epithelial apical surface with a solution of PBS (Phosphate Buffered Saline, Gibco) with 10 mM DTT (Dithiothreitol, ThermoFisher Scientific). To have a close-up view of the ciliary layer, the membrane supporting the ciliated epithelium was excised and removed from the plastic insert, carefully folded around a coverslip, and finally sandwiched between a microscope slide and another coverslip, creating a makeshift chamber (refer to (46) for more details). The chamber was filled with CMNB-Caged Fluorescein (Fluorescein bis-(5-Carboxymethoxy-2-Nitrobenzyl) Ether, Dipotassium Salt, ThermoFisher Scientific), diluted in a solution of 90% PBS and 10% culture medium (MucilAirTM culture medium, Epithelix S`arl) to a final concentration of 1-2 mM. As illustrated in Figure 1.A, a 12 ms pulse of 405 nm collimated light from a 200 mW laser diode (SLD3237VF, Sony), focused through the objective described below, was used to uncage the dye in a micrometric spot within the PCL. Photoactivation and imaging were carried out on a Nikon Eclipse Ti-E inverted microscope, using a 60× water immersion objective (NA = 1.2, Nikon) complemented by an additional 1.5× magnification such that 1 px = 0.0658 µm. A CMOS camera (GS3-U3-23S6M-C, FLIR Integrated Imaging Solutions Inc.) was used to capture videos at a rate of 80 fps (bright field) and 120-150 fps (epifluorescence). The fluorescent recording commenced approximately 0.30 s before the activation of the dye. For subsequent analyses, only frames post-dye activation were considered. This selection was determined by quantifying the average fluorescent intensity in each frame and considering only those subsequent to the frame exhibiting peak fluorescent signal. Typically, the peak signal frame exhibits saturation and is therefore omitted from analysis. Given that saturation occurs immediately post-activation, the first frame considered in the analysis is the second frame after dye activation, so it corresponds to a time point of ~ 0.016 s.

The radius of the activation spot was measured in ten fluorescent frames where the activation spot was visible without saturation of the fluorescent signal (similar to the one shown in Figure 1.C). To enhance spot visibility, the images were first binarised, followed by the application of a circle detection algorithm to identify and measure the radius of the circular region. Across all frames, the maximum measured radius was ~1.5 µm.

Image acquisition was managed by custom software on a Linux platform. Starting from the apical surface of the epithelium, the dye is activated in different positions, each 1.5 µm apart, progressing towards the ciliary tips. The custom software also allowed sample temperature regulation. Thermal stabilization was achieved by employing a glass heater adjacent to the chamber slide’s upper side and equipping the water immersion objective, which contacted the sample’s lower side, with a heating collar (see Figure 1.A).

### Video analysis

In our study, all video and data analysis are executed using scripts written MATLAB R2021a (The MathWorks Inc.).

#### Tracking of the fluorescent dye

Upon photo-activation of the caged dye, the fluid speed was measured as the displacement of the fluorescent patch along lines parallel to the ciliated epithelium. The fluorescent intensity profile is evaluated on a measuring strip orientated in the direction of the power stroke. The strip is found as the 20 pixel dilation of a measuring line obtained by manually tracing the cellular contour in the bright field video, and then translating this profile perpendicularly to the epithelium until it intersects with the dye activation point. The fluorescence intensity along the measuring strip was fitted with a Gaussian curve: 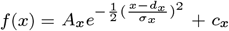. The mean *dx*(*t*) and width *σx*(*t*) of the Gaussian describe the displacement of the center of the fluorescent cloud and its spread along the measuring line over time, respectively. Example of a Gaussian fit applied to the dye fluorescence intensity profile on the measuring line for different frames after dye activation can be found in *SI Appendix*, Fig. 4.

We chose to explore the *z* component of the velocity field due to the consistent decrease in speed along *x* over time. In each frame following dye activation, a 500-pixel long vertical measuring line is positioned such that its *x* coordinate corresponds to the center of the dye cloud at that time, measured on the measuring line *dx*(*t*), while its midpoint aligns with the vertical coordinate of the measuring line. This segment runs parallel to the *z*-axis in each frame. Along the vertical line, the fluorescent data are fitted with a Gaussian curve to determine the vertical position of the center of the dye cloud: 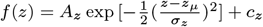. An example of this fit for four different times after dye activation can be found in *SI Appendix*, Fig. 5. The vertical position of the cloud is normalised to the position of the epithelium, calculated as the distance between the vertical position of the Gaussian mean *zµ*(*t*) and the epithelium *z*_*epithelium*_(*t*). This yields the position of the dye along the *z* axis relative to the epithelium, denoted as *dz* (*t*).

#### Measurement of dye diffusion

Prior to experiments, three highspeed fluorescent videos were collected in the area of the chamber farthest from the cilia at room temperature. We specifically chose this location to closely observe how the dye spreads following photoactivation, ensuring that cilia movement did not interfere with our observations. The fluorescent profile of the dye was evaluated over time on a 1000 pixel horizontal line passing through the activation spot using the algorithm described above: a Gaussian function was fitted to the data to measure the standard deviation of this distribution over time, providing insights into the spread of the dye (more info and data in *SI Appendix*, section 3.C). We approached this problem as a diffusion process from an instantaneous point source, which is well-described by an analytical solution (70). For the initial 0.2 seconds after dye activation, the probability density of the distribution was fitted with the equation 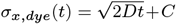. Here, *D* and *C* are parameters. *D* represents the dye diffusion coefficient, and *C* an intercept term added to account for the finite size of the source, influenced by the activation laser. From fits of the standard deviation across three experimental runs at room temperature, we determined 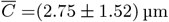 µm and 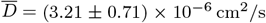. The average size of the dye source matches with the size of the initial dye spot measured in the videos (maximum 1.5 µm radius), as described previously. The estimated value of *D* aligns well with previously reported diffusion coefficients for fluorescein at 20 °C, as detailed in (71, 72). This agreement supports the methods and analysis we used in our experiments.

For the experiments focusing on dye activation in the ciliary layer, we adjusted our methodology. We introduced a modification to the previous equation to model the standard deviation of the Gaussian fit for the first 0.15 seconds after dye activation: 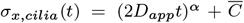. In this model *α* and *D*_*app*_ are newly introduced parameters that account for the altered diffusion dynamics in the cilia layer, 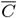 is the average size of the dye source measured as described above. Examples of this fit can be found in Figure 2.B where *σx* is plotted on a logarithmic scale for different dye activation positions.

#### CBF assessment

In every bright field video, the *CBF* was measured at three distinct locations within the ciliary layer, horizontally spaced 150 pixels (~10 µm) apart and situated 70 pixels (~4.5 µm) away from the epithelium contour. The vertical position was selected to be roughly in the middle of the cilia layer, between the cilia base and tips. At each of these positions, *CBF* was estimated in concentric boxes of 40, 60 and 80 pixels by side. For any given box *B*_*i*_, the dominant frequency was identified as the location of the highest peak of the first harmonic of the average periodogram, 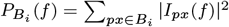 the FFT. Subsequently, the *CBF* for each of the three reference points was determined as the mean of the frequencies in the three boxes. The overall representative *CBF* for the video was then computed as the average *CBF* across these three locations (more info in *SI Appendix*, section 1).

#### Kymograph Generation and 2D Autocorrelation

As reported in (51, 73, 74), ciliary synchronizations patterns can be detected using kymographs. Grey-scale intensity kymographs are generated on 100 pixel long segments. This segments are roughly in the middle of the ciliary layer, parallel to the epithelium and orientated in the same direction of the power stroke. Grey-scale intensities measured from each segment were cross-correlated to determine the spatio-temporal coordination exhibited by the beating cilia.

## Data, Materials, and Software Availability

Data analysis codes are available at https://github.com/erikacausa.

## ACKNOWLEDGMENTS

The authors thank Maurizio Chioccioli for preliminary experiments, and Morten Kals for useful assistance and insights during the experimental setup and planning phase. This study was supported by the European Union’s Horizon 2020 research and innovation program under the Marie Sk-lodowskaCurie grant agreement No 955910 (ITN PHYMOT) and the UK CF Trust SRC016.

## References

1. M Legendre, LE Zaragosi, HM Mitchison, Motile cilia and airway disease. Semin. cell & developmental biology 110, 19–33 (2021).

2. XM Bustamante-Marin, LE Ostrowski, Cilia and mucociliary clearance. Cold Spring Harb. perspectives biology 9 (2017).

3. P Satir, MA Sleigh, The physiology of cilia and mucociliary interactions. Annu. review physiology 52, 137–155 (1990).

4. JV Fahy, BF Dickey, Airway mucus function and dysfunction. New Engl. journal medicine 363, 2233–2247 (2010).

5. AE Tilley, MS Walters, R Shaykhiev, RG Crystal, Cilia dysfunction in lung disease. Annu. review physiology 77, 379–406 (2015).

6. A Wanner, M Salathé, TG O’Riordan, Mucociliary clearance in the airways. Am. J. Respir. Critical Care Medicine 154, 1868–1902 (1996) PMID: 8970383.

7. MR Knowles, RC Boucher, et al., Mucus clearance as a primary innate defense mechanism for mammalian airways. The J. clinical investigation 109, 571–577 (2002).

8. JC Nawroth, AM van der Does, A Ryan Firth, E Kanso, Multiscale mechanics of mucociliary clearance in the lung. Philos. transactions Royal Soc. London. Ser. B, Biol. sciences 375, 20190160 (2020).

9. MR Knowles, LA Daniels, SD Davis, MA Zariwala, MW Leigh, Primary ciliary dyskinesia. recent advances in diagnostics, genetics, and characterization of clinical disease. Am. journal respiratory critical care medicine 188, 913–922 (2013).

10. A Livraghi, SH Randell, Cystic fibrosis and other respiratory diseases of impaired mucus clearance. Toxicol. pathology 35, 116–129 (2007).

11. M King, Physiology of mucus clearance. Paediatr. respiratory reviews 7, S212–S214 (2006).

12. E Puchelle, J Zahm, D Quemada, Rheological properties controlling mucociliary frequency and respiratory mucus transport. Biorheology 24, 557–563 (1987).

13. P Cicuta, The use of biophysical approaches to understand ciliary beating. Biochem. Soc. Transactions 48, 221–229 (2020).

14. DJ Thornton, JK Sheehan, From mucins to mucus: toward a more coherent understanding of this essential barrier. Proc. Am. Thorac. Soc. 1, 54–61 (2004).

15. SM Kreda, CW Davis, MC Rose, Cftr, mucins, and mucus obstruction in cystic fibrosis. Cold Spring Harb. perspectives medicine 2 (2012).

16. BC Huck, et al., Macro- and microrheological properties of mucus surrogates in comparison to native intestinal and pulmonary mucus. Biomacromolecules 20, 3504–3512 (2019).

17. DB Hill, B Button, M Rubinstein, RC Boucher, Physiology and pathophysiology of human airway mucus. Physiol. Rev. 102, 1757–1836 (2022).

18. W Hofmann, B Asgharian, The effect of lung structure on mucociliary clearance and particle retention in human and rat lungs. Toxicol. Sci. 73, 448–456 (2003).

19. SH Randell, RC Boucher, Effective mucus clearance is essential for respiratory health. Am. journal respiratory cell molecular biology 35, 20–28 (2006).

20. E Loiseau, et al., Active mucus–cilia hydrodynamic coupling drives self-organization of human bronchial epithelium. Nat. Phys. 16, 1158–1164 (2020).

21. SK Lai, YY Wang, D Wirtz, J Hanes, Micro-and macrorheology of mucus. Adv. drug delivery reviews 61, 86–100 (2009).

22. M Jory, et al., Mucus from human bronchial epithelial cultures: rheology and adhesion across length scales. Interface Focus. 12, 20220028 (2022).

23. M Braunreuther, M Liegeois, JV Fahy, GG Fuller, Nondestructive rheological measurements of biomaterials with a magnetic microwire rheometer. J. Rheol. 67, 579–588 (2023).

24. J Widdicombe, Regulation of the depth and composition of airway surface liquid. J. anatomy 201, 313–318 (2002).

25. B Button, et al., A periciliary brush promotes the lung health by separating the mucus layer from airway epithelia. Sci. (New York, N.Y.) 337, 937–941 (2012).

26. H Hendrik, E Raubenheimer, The role of airway surface liquid in the primary management of rhinosinusitis. J Interdiscipl Med Dent Sci 1, 2 (2013).

27. H Matsui, SH Randell, SW Peretti, CW Davis, RC Boucher, Coordinated clearance of periciliary liquid and mucus from airway surfaces. The J. clinical investigation 102, 1125–1131 (1998).

28. J Hussong, et al., Cilia-driven particle and fluid transport over mucus-free mice tracheae. J. biomechanics 46, 593–598 (2013).

29. S Bermbach, et al., Mechanisms of cilia-driven transport in the airways in the absence of mucus. Am. journal respiratory cell molecular biology 51, 56–67 (2014).

30. JR Blake, A spherical envelope approach to ciliary propulsion. J. Fluid Mech. 46, 199–208 (1971).

31. F Boselli, J Jullien, E Lauga, RE Goldstein, Fluid mechanics of mosaic ciliated tissues. Phys. Rev. Lett. 127, 198102 (2021).

32. GR Ramirez-San Juan, et al., Multi-scale spatial heterogeneity enhances particle clearance in airway ciliary arrays. Nat. Phys. 16, 958–964 (2020).

33. J Han, CS Peskin, Spontaneous oscillation and fluid–structure interaction of cilia. Proc. Natl. Acad. Sci. 115, 4417–4422 (2018).

34. J Elgeti, G Gompper, Emergence of metachronal waves in cilia arrays. Proc. Natl. Acad. Sci. 110, 4470–4475 (2013).

35. DJ Smith, A boundary element regularized stokeslet method applied to cilia-and flagella-driven flow. Proc. Royal Soc. A: Math. Phys. Eng. Sci. 465, 3605–3626 (2009).

36. DJ Smith, EA Gaffney, JR Blake, Mathematical modelling of cilia-driven transport of biological fluids. Proc. R. Soc. Lond. A 465, 2417–2439 (2009).

37. T Niedermayer, B Eckhardt, P Lenz, Synchronization, phase locking, and metachronal wave formation in ciliary chains. Chaos: An Interdiscip. J. Nonlinear Sci. 18 (2008).

38. N Uchida, R Golestanian, Generic conditions for hydrodynamic synchronization. Phys. Rev. Lett. 106, 058104 (2011).

39. A Maestro, et al., Control of synchronization in models of hydrodynamically coupled motile cilia. Commun. Phys. 1, 058104 (2018).

40. AV Kanale, F Ling, H Guo, S Fürthauer, E Kanso, Spontaneous phase coordination and fluid pumping in model ciliary carpets. Proc. Natl. Acad. Sci. 119, e2214413119 (2022).

41. Wh Li, G Zheng, Photoactivatable fluorophores and techniques for biological imaging applications. Photochem. & photobiological sciences : Off. journal Eur. Photochem. Assoc. Eur. Soc. for Photobiol. 11, 460–471 (2012).

42. WR Lempert, H Scott Raymond, Flow tagging velocimetry using caged dye photo-activated fluorophores. Meas. Sci. Technol. pp. 1251–1258 (2000).

43. JP Shelby, DT Chiu, Mapping fast flows over micrometer-length scales using flow-tagging velocimetry and single-molecule detection. Anal. Chem. 75, 1387–1392 (2003).

44. W Lempert, P Ronney, K Magee, K Gee, R Haugland, Flow tagging velocimetry in incompressible flow using photo-activated nonintrusive tracking of molecular motion (phantomm). Exp. Fluids 18, 249–257 (1995).

45. GC Ellis-Davies, Caged compounds: photorelease technology for control of cellular chemistry and physiology. Nat. methods 4, 619–628 (2007).

46. E Causa, R Fradique, P Cicuta, Measuring biophysical properties of cilia motility from mammalian tissues via quantitative video analysis methods in Cilia: Methods and Protocols. (Springer), pp. 251–262 (2023).

47. A Oltean, AJ Schaffer, PV Bayly, SL Brody, Quantifying ciliary dynamics during assembly reveals stepwise waveform maturation in airway cells. Am. journal respiratory cell molecular biology 59, 511–522 (2018).

48. R Williams, N Rankin, T Smith, D Galler, P Seakins, Relationship between the humidity and temperature of inspired gas and the function of the airway mucosa. Critical care medicine 24, 1920–1929 (1996).

49. AD Workman, NA Cohen, The effect of drugs and other compounds on the ciliary beat frequency of human respiratory epithelium. Am. journal rhinology & allergy 28, 454–464 (2014).

50. PG Jayathilake, Z Tan, DV Le, HP Lee, BC Khoo, Three-dimensional numerical simulations of human pulmonary cilia in the periciliary liquid layer by the immersed boundary method. Comput. & Fluids 67, 130–137 (2012).

51. KY Wan, et al., Reorganization of complex ciliary flows around regenerating stentor coeruleus. Philos. Transactions Royal Soc. B 375, 20190167 (2020).

52. S Chateau, U D’Ortona, S Poncet, J Favier, Transport and mixing induced by beating cilia in human airways. Front. physiology 9, 161 (2018).

53. Y Ding, JC Nawroth, MJ McFall-Ngai, E Kanso, Mixing and transport by ciliary carpets: a numerical study. J. Fluid Mech. 743, 124–140 (2014).

54. A Burn, et al., A quantitative interspecies comparison of the respiratory mucociliary clearance mechanism. Eur. biophysics journal 51, 51–65 (2022).

55. L Feriani, et al., Assessing the collective dynamics of motile cilia in cultures of human airway cells by multiscale ddm. Biophys. journal 113, 109–119 (2017).

56. J Kotar, M Leoni, B Bassetti, M Cosentino Lagomarsino, P Cicuta, Hydrodynamic synchronization of colloidal oscillators. Proc. Natl. Acad. Sci. USA 107, 7669–7673 (2010).

57. C Pozrikidis, Boundary integral and singularity methods for linearized viscous flow. (Cambridge university press), (1992).

58. S Kim, SJ Karrila, Microhydrodynamics: principles and selected applications. (Butterworth-Heinemann), (2013).

59. J Happel, H Brenner, Low Reynolds number hydrodynamics: with special applications to particulate media. (Springer Science & Business Media) Vol. 1, (2012).

60. E Lauga, The fluid dynamics of cell motility. (Cambridge University Press) Vol. 62, (2020).

61. J Gray, GJ Hancock, The propulsion of sea-urchin spermatozoa. J. Expt. Biol. 32, 802–814 (1955).

62. J Lighthill, Flagellar hydrodynamics. SIAM review 18, 161–230 (1976).

63. RE Johnson, An improved slender-body theory for stokes flow. J. Fluid Mech. 99, 411–431 (1980).

64. GR Fulford, JR Blake, Muco-ciliary transport in the lung. J. Theo. Biol. 121, 381–402 (1986).

65. E Lauga, TR Powers, The hydrodynamics of swimming microorganisms. Rep. Prog. Phys. 72, 096601 (2009).

66. N Pellicciotta, et al., Cilia density and flow velocity affect alignment of motile cilia from brain cells. J. Expt. Biol. 223, jeb229310 (2020).

67. J Devalia, R Sapsford, C Wells, P Richman, R Davies, Culture and comparison of human bronchial and nasal epithelial cells in vitro. Respir. medicine 84, 303–312 (1990).

68. Y Fu, et al., Ciliostasis of airway epithelial cells facilitates influenza a virus infection. Vet. research 49, 1–4 (2018).

69. R Robinot, et al., Sars-cov-2 infection induces the dedifferentiation of multiciliated cells and impairs mucociliary clearance. Nat. communications 12, 4354 (2021).

70. J Crank, The mathematics of diffusion. (Oxford university press), (1979).

71. C Soeller, et al., Application of two-photon flash photolysis to reveal intercellular communication and intracellular ca 2+ movements. J. biomedical optics 8, 418–427 (2003).

72. CT Culbertson, SC Jacobson, JM Ramsey, Diffusion coefficient measurements in microfluidic devices. Talanta 56, 365–373 (2002).

73. C Ringers, et al., Novel analytical tools reveal that local synchronization of cilia coincides with tissue-scale metachronal waves in zebrafish multiciliated epithelia. Elife 12, e77701 (2023).

74. DR Brumley, M Polin, TJ Pedley, RE Goldstein, Metachronal waves in the flagellar beating of volvox and their hydrodynamic origin. J. Royal Soc. Interface 12, 20141358 (2015).

