## Supplementary material for "Cilia dynamics creates a dynamic barrier to penetration of the periciliary layer in human airway epithelia"

### 1. CBF assessment

**A. Measurement in concentric boxes.** In each video, the cilia beating frequency (CBF) was quantified at three specific reference points within the ciliary layer. These points were horizontally spaced 150 pixels apart and located 70 pixels from the epithelium contour. At each reference point, $CBF$ was estimated using concentric boxes sized at 40, 60, and 80 pixels per side (as depicted in figure S1.A). The center of each box was positioned at approximately half the length of the cilia, consistent across various dye activation positions. For each pixel within a box, the frequency was identified at the peak of the first harmonic in the intensity periodogram. The characteristic frequency for the box was calculated as the average of the frequency distribution for all pixels in the box, with measurement error represented by the standard deviation of this distribution. As shown in figure S1.B, there were variations in the measured characteristic frequency across different reference locations, but these did not significantly vary with the size of the measuring box. The $CBF$ at each reference point was determined by averaging the frequencies from the three boxes.

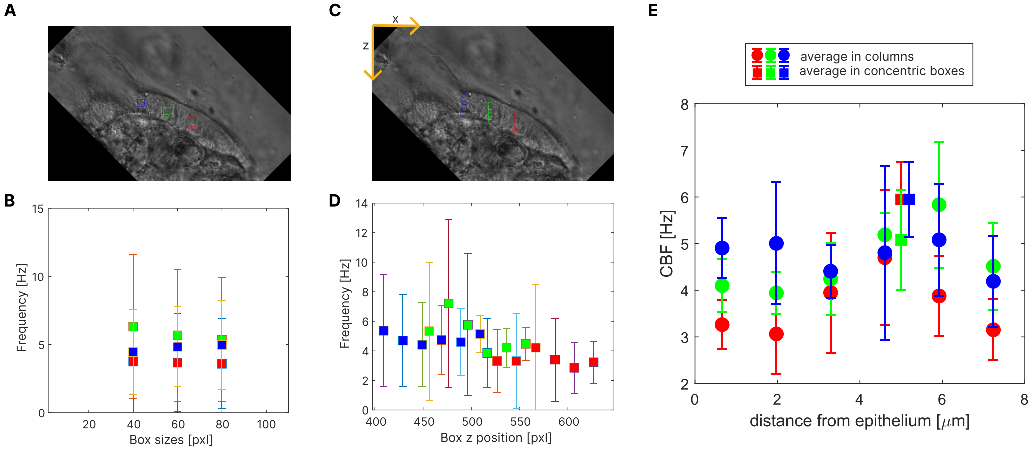


**Fig. S1.** Measurement of $CBF$ in concentric boxes (A,B) and columnar boxes (C,D). (E) Measurement of $CBF$ in columns for different fields of view of the same region (different dye activation spot). Data in figure is from experiment carried out at room temperature. Scale bars are 10 $\mu m$.

**B. Measurement in vertical columns.** We validated that the CBF measurement is not affected by the vertical position of the measuring box's center. Specifically, we measured the characteristic frequency in six boxes, each 20 pixels by side. These boxes shared the same $x$-coordinate as the center of the previously used concentric boxes, but their $z$-coordinate varied from the base to the tip of the cilium, creating a columnar arrangement (figure S1.C). The characteristic frequency for each box was measured as previously described and then plotted against the $z$-coordinate of the box's center (shown in figure S1.D). This plot revealed that the mean frequency does not significantly change with the box's position. Further, the average $CBF$ at each reference $x$-coordinate was computed by averaging the frequencies from the three boxes. In figure S1, we depicted the average $CBF$ for the three reference points across different fields of view, corresponding to various dye activation spots, using circles. Notably, at position 4, $CBF$ was also measured using concentric boxes (represented by squares in the figure). The congruence of these data, within error margins, supports our conclusion: the method of measurement, whether using columnar or concentric boxes, does not significantly impact the $CBF$ results. Additionally, it is evident that the $z$-coordinate of the box is not a critical factor, provided that the box is entirely within the ciliary layer.

**C. CBF with repeated dye activation.** As detailed in the main manuscript, the dye was activated at 5-7 different positions in the ciliary layer, progressing from the epithelium towards the tips of the cilia. To assess whether multiple laser exposures would influence ciliary beating, we investigated the impact of repetitive dye activation on cilia. For this purpose, we measured $CBF$ using concentric boxes across all experiments conducted at various positions. The $CBF$ data were normalized by dividing the measured value by the $CBF$ measured at the first activation position $z_{min}$ and averaged in groups depending on the positions of the dye activation within the ciliary layer. Finally, these normalized $CBF$ values are plotted against dye activation height $z$. Results at RT and 37$^o$C can be found in figure S2 A and B, respectively. We observed an average reduction in $CBF$ of about 15% in both cases, meaning that repeated laser activation affect ciliary beating. However, while this reduction is statistically significant, it is relatively modest in magnitude.

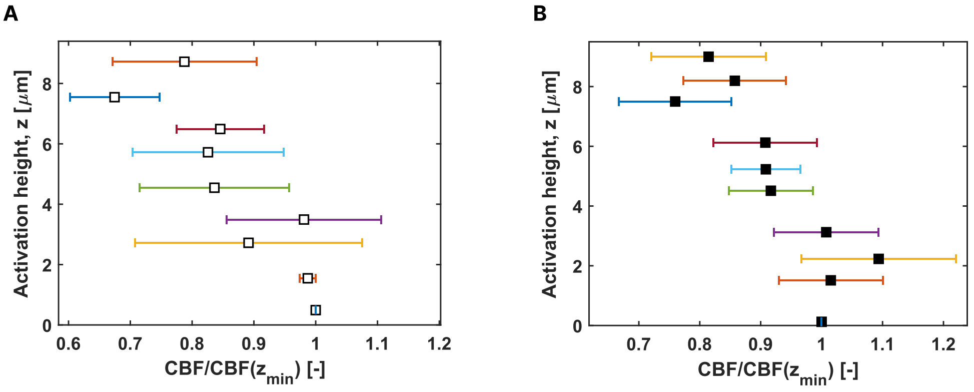


**Fig. S2.** Analysis of CBF relative to dye activation locations in the ciliary layer. We investigated the impact of repetitive dye activation on $CBF$ measurement. Data is from (A) measurements conducted at room temperature, and (B) measurements performed at a controlled temperature of 37$^o$C.

### 2. Measurement of cilium speed

In instances where individual cilia could be distinctly observed throughout their entire beating pattern, we measured the instantaneous velocity of the cilium. This was achieved by selecting points in 20-30 consecutive frames where the cilium intersected with lines positioned 1, 2, and 3 $\mu m$ away from the cilium base and aligned parallel to the direction of beating (as shown by dashed lines in Figure S3.A). We then quantified the mean displacement of the cilium between consecutive frames for both the power and recovery strokes. This was done by measuring the mean square displacement between the collected coordinates (Figure S3.B), enabling us to calculate the average speed of the cilium during the two distinct phases of its beating cycle. Subsequently, the net speed of the cilium was determined by calculating the difference between the speeds during the power and recovery strokes, as indicated by green marks in Figure S3.C. From three repeated measurements conducted at room temperature, the average cilium speed was found to be $18.3 \pm 4.2\,(\mu m/s)$, which is approximately twice the average fluid velocity recorded over short-time intervals $10.5 \pm 7.6\,(\mu m/s)$ and nearly six times the average velocity over long-time intervals $3.0 \pm 1.1\,(\mu m/s)$.

### 3. Dye displacement and diffusion

**A. Displacement and diffusion along lines parallel to epithelium (*x*).** Upon photo-activation of the caged dye, the fluid speed was measured as the displacement of the fluorescent patch along lines parallel to the ciliated epithelium. The fluorescent intensity profile is evaluated on a measuring strip orientated in the direction of the power stroke (white line and arrow in figure S4.A). The strip is found as the 20 pixel dilation of a measuring line obtained by manually tracing the cellular contour in the bright field (BF) video, and then translating this profile perpendicularly to the epithelium until it intersects with the dye activation point (red cross in figure S4.A). The purpose of the dilation is to achieve better statistics, as the quantities that we measure in a point will be averaged over the small region of space perpendicular to the measuring line at that point. The user orients the measuring strip so that its arclength increases in the direction of the power stroke. In each frame after the activation of the dye we measure the fluorescence intensity along the measuring strip and fit it with a Gaussian curve:

$$f(x) = A_x e^{-\frac{1}{2}(\frac{x-d_x}{\sigma_x})^2} + c_x \qquad [S1]$$

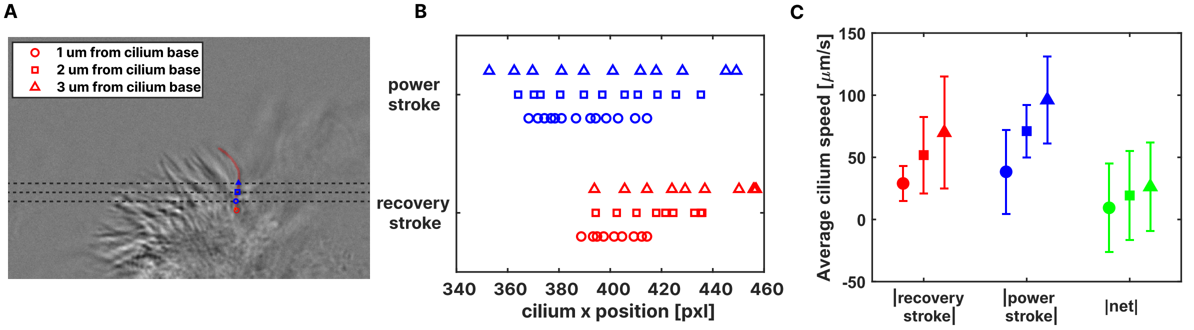


**Fig. S3.** (A) Cilium speed is measured by tracking the motion of a selected cilium (in red) at its intersection with lines parallel to the direction of the power strokes and distant 1,2 and 3 $\mu m$ from the cilium base (red spot). (B) The x-coordinates of the cilium at different distances are stored to measure the mean displacement during power and recovery stroke. (C) The average speed for the two strokes are measured and the net cilium speed calculated as the difference between the two speeds. All data were gathered from experiments conducted at room temperature.

where $d_x(t)$ and $\sigma_x(t)$ represented the mean and width of the Gaussian curve, respectively. These parameters yielded information about the displacement of the center of the fluorescent patch (Figure S4.B) and its spread along the measuring line over time (Figure S4.C), providing insights into fluid velocity and dye diffusion, respectively.

Example of a Gaussian fit applied to the dye fluorescence intensity profile on the measuring line for different frames after dye activation can be found in figure S4.D. Notably, the fluorescence intensity distribution exhibits asymmetry. This deviation from symmetry is attributed to the lateral displacement of the dye's center of mass induced by the fluid flow dynamics generated by ciliary motion. Despite this asymmetry, a standard Gaussian fit proves to be effective in closely determining both the central location and the extent of the fluorescent patch.

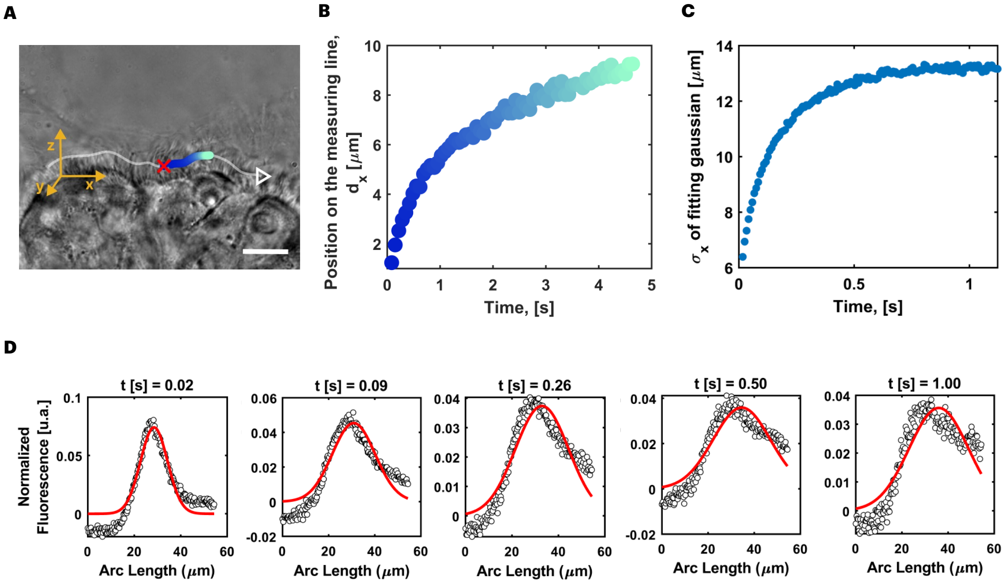


**Fig. S4.** (A) The fluorescent intensity profile is measured along a measuring strip (indicated by the white line). The strip is orientated in the cilia's power stroke direction and passes through the dye activation spot (red cross). (D) For each frame, the fluorescence intensity along the strip is well fitted by a Gaussian curve. The experimental data are depicted as black circles, while the fit, represented by equation S1, is shown in red. (B) The displacement of the mean of the fluorescent cloud quantifies the transport (blue to light-blue circles), while (C) its spread (blue small circles) provides information about the dye diffusion. Scale bar = 10 $\mu m$.

**B. Displacement and diffusion along lines perpendicular to the epithelium (*z*).** We chose to explore the $z$ component of the velocity field due to the consistent decrease in speed along $x$ over time. In each epi-fluorescence (EPI) frame following dye activation, a 500-pixel long vertical measuring line is positioned such that its $x$ coordinate corresponds to the center of the dye cloud at that time, measured on the measuring line ($d_x(t)$), while its midpoint aligns with the vertical coordinate of the measuring line. This segment runs parallel to the $z$-axis in each frame. Along the vertical line, the fluorescent data are fitted with a Gaussian curve to determine the vertical position of the center of the dye cloud:

$$f(z) = A_z e^{-\frac{1}{2}(\frac{z-z_\mu}{\sigma_z})^2} + c_z \quad [S2]$$

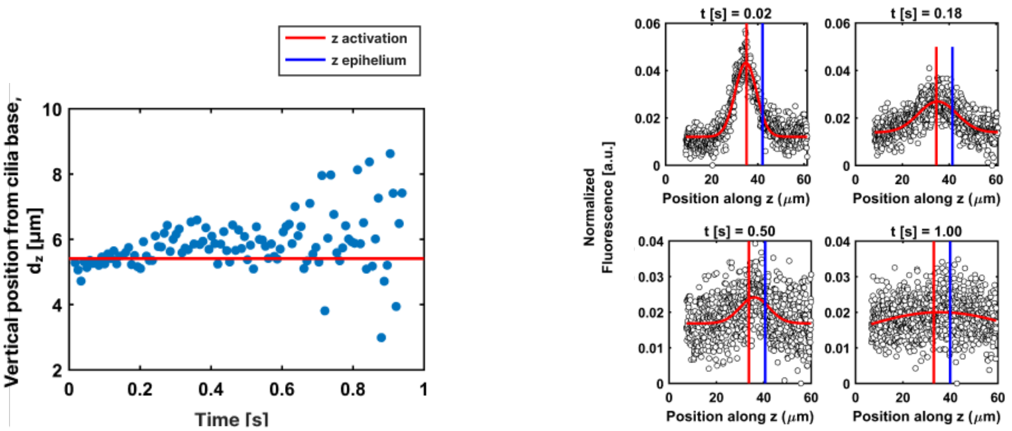


**Fig. S5. Dye displacement along $z$.** The displacement of the centre of the fluorescent cloud (A) is measured along lines perpendicular to the epithelium by fitting the fluorescent data (circle markers) with a Gaussian curve (in red) over time, as in equation S2 (B). The red line indicates the vertical coordinate at which the dye is activated, while the blue line indicates the epithelium position.

An example of this fit for four different times after dye activation can be seen in Figure S5.B, where the data points are represented by circle markers and the fit is depicted in red. In this figure, the red and blue lines specifically indicate the positions where the dye is released $z_{dye}$ and the location of the epithelium $z_{epithelium}$ respectively.

The vertical position of the cloud is normalised to the position of the epithelium, calculated as the distance between the vertical position of the Gaussian mean $z_{\mu}(t)$ and the epithelium $z_{epithelium}(t)$. This yields the position of the dye along the $z$ axis relative to the epithelium, denoted as $d_z(t)$. A typical dye plot of dye displacement along $z$ is presented in figure S5.A. In this plot, after activation, the dye moves towards the tip of the cilia.

**C. Dye diffusion in watery liquid.** Three high-speed fluorescent videos were collected in the area of the chamber farthest from the cilia at RT. We analysed the fluorescent profile of the dye over time along a 1000-pixel long horizontal line that intersected the dye activation spot (depicted as a blue line in Figure S6.A). To enhance statistical reliability, this line was dilated by 20 pixels. As illustrated in Figure S6.D, we fitted a Gaussian function (in red) to the data to measure the mean and the standard deviation of this distribution over time, as described previously by equation S1. Plots of these quantities can be found in Figure S6.B and C, respectively. The spread of the data in the displacement plot can be attributed to a decrease in the algorithm's accuracy when the peak intensity of the dye cloud becomes less pronounced (see last panel in Figure S6.D).

We approached this problem as a diffusion process from an instantaneous point source, which is well-described by an analytical solution. For the initial 0.2 seconds after dye activation, the probability density of the distribution was fitted with the equation:

$$\sigma_{x,dye}(t) = \sqrt{2Dt} + C \qquad [S3]$$

Here, $D$ and $C$ are parameters. $D$ represents the dye diffusion coefficient, and $C$ an intercept term added to account for the finite size of the source, influenced by the activation laser.

**D. Linear dye displacement.** In a subset of the experiments, the dye displacement over time exhibited a linear pattern, observed in 13% of the cases at $37^{o}$C and in 66% at room temperature (RT). For these specific instances, such as the 3rd and 5th activation positions illustrated in Figure S7.B, dye displacement along $x$ is linear, correlated with a constant speed measured as the slope of the line fitting the data, as depicted by the black dotted line in Figure S7.C. In all cases, we observed a tendency of the cilia to push the activated dye patch towards the tips when this is activated within the ciliary layer.

### 4. PCL fluid flow at room temperature

We investigated the influence of temperature on PCL transport by repeating the transport experiments at room temperature. Specifically we conducted 9 experiments with 2 distinct inserts where the sample temperature ranged between 22–24 °C. Notably, the short-time and long-time transport at RT displayed trends consistent with those observed at the physiological temperature of 37 °C (Figure S8.A). However, a higher incidence of linear transport patterns was observed at RT, occurring in 66% of these experiments. Figure S8.B presents the comparative results of experiments conducted at the two temperatures, illustrating that a reduction in temperature from $37^{o}$C to room temperature results in a 20.9% decrease in $CBF$. This reduction has a pronounced effect on dye clearance speed, with the short-time speed decreasing by 44.1% and long-time speed decreasing by 21.9% reduction. These findings underscore the susceptibility of mucociliary transport to temperature fluctuations and confirm a strong correlation between short-time flow speed and $CBF$.

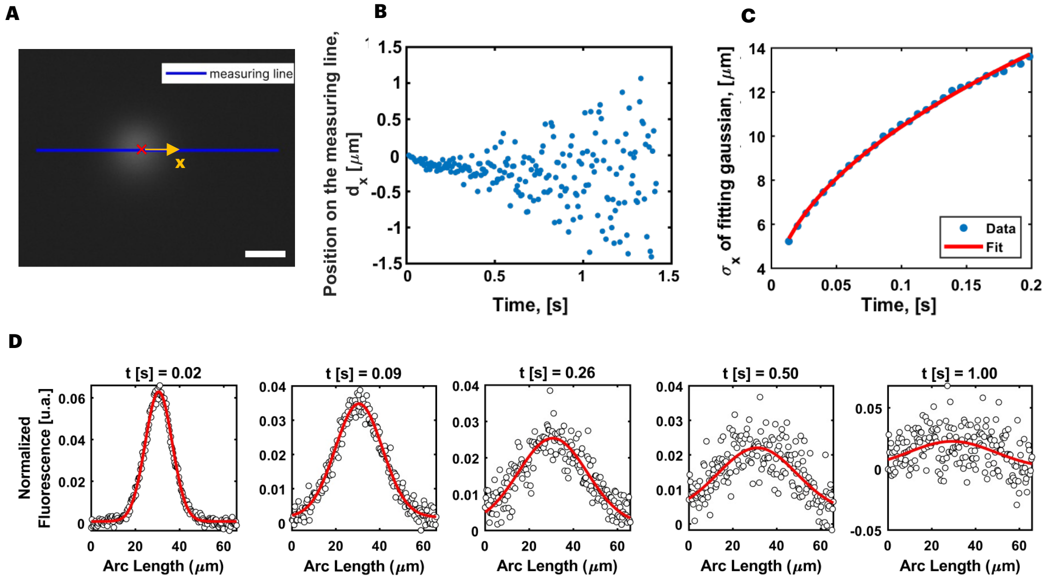


**Fig. S6.** The caged dye in activated far from the ciliary layer at room temperature. The fluorescent profile is fitted on a measuring segment (A) with a Gaussian curve (fit in red) (D) over time, as described in eq. S1. (B) In the absence of beating cilia, the center of the dye patch remains stationary. (C) Up until $t = 0.2$ seconds, the dye's spread follows the diffusion equation from a instantaneous point source (eq. S3), allowing the calculation of the dye's diffusion coefficient based on this fit (in red). Scale bar = 10 $\mu m$.

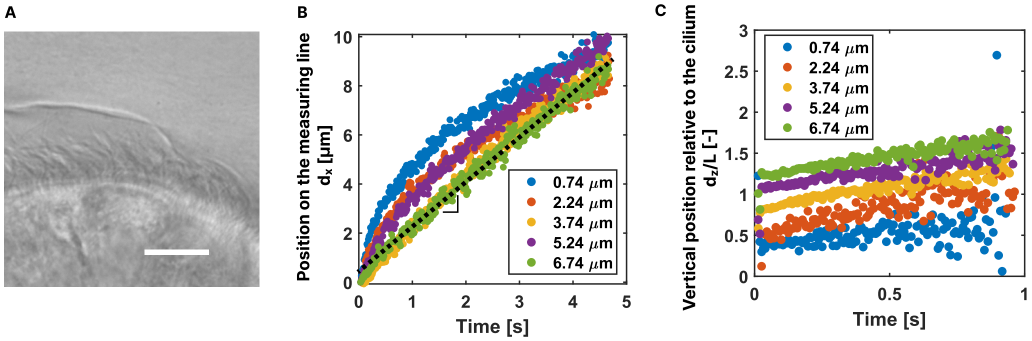


**Fig. S7.** In some experiments, as the one presented in (A), a linear displacement of the dye along the $x$-axis is observed, particularly at specific dye activation positions (in this case the 3rd and 5th positions in B). This is associated also to constant speed along $z$ (C). Also in this experiments the dye is pushed outside the PCL, towards the cilia tips. Scale bar = 10 $\mu m$.

### 5. Correlation between PCL short-time speed and CBF

To explore the correlation between long-time speed and CBF, we considered a non-linear model to capture the potential peak in speed at an optimal CBF, above which the speed decreases. We propose an empirical model:

$$U_{x,long}(CBF) = \frac{p_1 CBF}{p_2 + CBF^2} \qquad [S4]$$

This model hypothesises an 'optimal' CBF for maximum long-time speed, after which the speed declines. Similarly to the previous fit, this model also predicts zero speed at zero CBF. The measured parameters for the fits, shown by lines in figure S9, are for $z$<4 $\mu m$ $p_1 = 23.6$ $\mu m/\mathrm{s}^3$ and $p_2 = 26.2$ $1/\mathrm{s}^2$, and for 4 $\mu m \leq z$<7 $\mu m$ $p_1 = 25.4$ $\mu m/\mathrm{s}^3$ and $p_2 = 10.7$ $1/\mathrm{s}^2$. Although the model did not perfectly fit the data, the discrepancies could be attributed to the limited data available. Nonetheless, this finding opens up a promising area for future research, inviting a deeper exploration into the relationship between CBF and optimal PCL speed.

### 6. Oscillation curve fitting

In instances where oscillatory behavior was noted in the dye displacement plot, the data was fitted using a combination of a sinusoidal $f(t) = A\sin(Bt + C)$ and a polynomial function $g(t) = Dt + Et^2 + Ft^3$, as illustrated in figure S10. For each curve the amplitude and period have been measured as $A$ and $2\pi/B$, respectively.

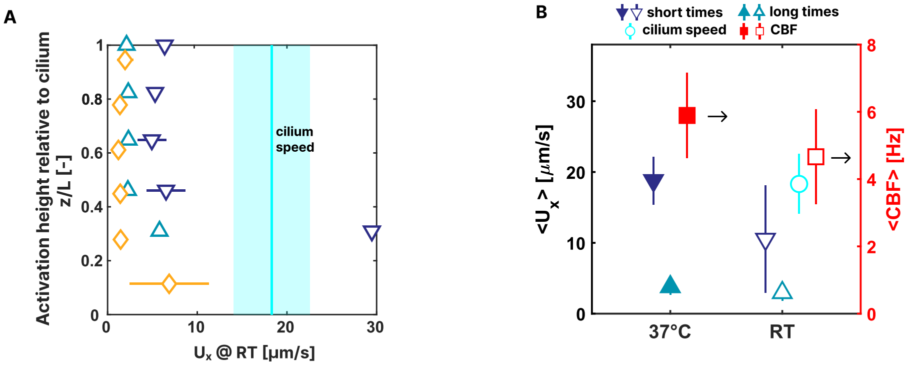


**Fig. S8.** (A) The PCL average velocity profile for experiments conducted at room temperature is presented here. The average length of cilia was measured to be 8.7 $\pm$ 0.2 $\mu m$. Both short-time and long-time speeds follow a pattern akin to that observed at 37℃, with the occurrence of linear behavior being more prevalent, accounting for 66% of the experiments. Anomalies observed in the data for low height in the PCL may be attributable to insufficient data points due to difficulties in shining the laser close to the apical surface of the epithelium. (B) Summary of main results from experiments conducted at physiological temperature and RT. As temperature declines from 37℃ to RT, $CBF$ experiences a 20.9% drop and flow speed decreases drastically (short-time speed reduction is $>$40%, while long-time speed $>$20%

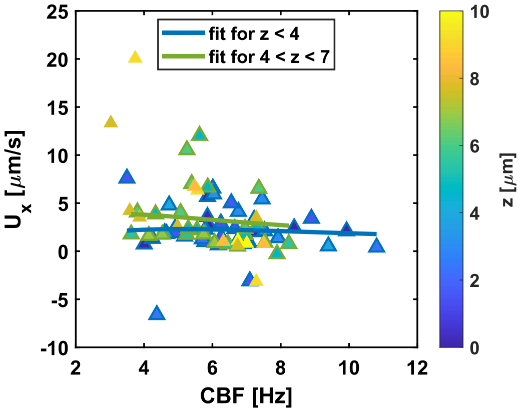


**Fig. S9.** Long-time speed data are grouped by the $z$-axis locations of dye activation and fitted with the nonlinear model described in equation S4 which hypothesises an 'optimal' CBF for maximum long-time speed, after which the speed declines.

### 7. Analysis of the autocorrelation plots

**Movie S1.** Video of ciliary patches displaying the synchronization patterns shown in main document, fig. 4. The movies are played at 0.5$\times$ speed, with scale bars representing 10 $\mu m$. The top left panel shows asynchronous cilia, while the bottom left panel shows fully synchronized cilia. The top right panel depicts a symplectic metachronal wave, and the bottom right panel shows an antiplectic MW.

**A. Metachronal wave speed.** We focused on two experiments where a phase lag between neighbouring cilia has been detected to measure the MW speed. The first one, whose autocorrelation plot is shown in figure main document, 4.I presents a symplectic MW whilst the second, in figure main document, 4.L, presents an antiplectic MW. We examined the behavior of autocorrelation against lag time on a 100-pixel segment at a constant space shift $S$ (specifically for shifts of 10, 30, and 50 pixels in the symplectic MW example proposed in figure S11.A,D and for shifts of 10, 20, and 30 pixels for the antiplectic MW in figure S12.E,F). The data were fitted using a damped oscillator equation:

$$A(\delta t) = H\cos(\omega\delta t + \Phi)e^{-\gamma\delta t} + K \quad \text{[S5]}$$

With $H$, $\omega$, $\Phi$, $\gamma$, and $K$ parameters.

We measured the MW speed as:

$$U_{MW} = <\Delta S/\Delta\Phi> \quad \text{[S6]}$$

The speed of MW propagation measured was $0.79 \pm 0.18$ mm/s for the symplectic wave and $0.17 \pm 0.08$ mm/s for the antiplectic wave.

For the scenario of in-phase cilia beating, we again explored the behavior of autocorrelation over a 100-pixel segment, examining its relationship with lag-time at constant spatial shifts of 25, 50, and 75 pixels (figure S12.A-C). The data were

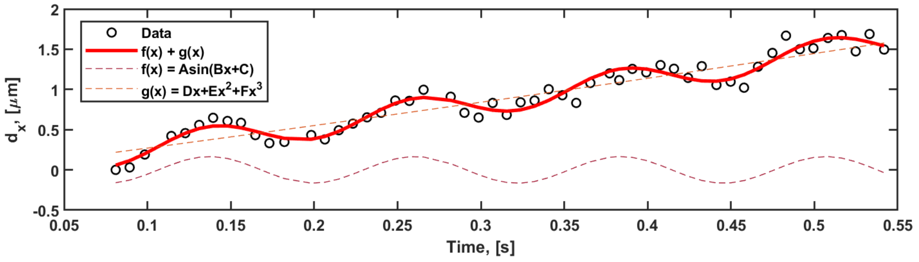


**Fig. S10.** Emergence of oscillatory fluid flow at short timescales is seen in some experiments. In the specific time interval where the dye displacement exhibits oscillatory behavior, a sinusoidal curve fitting is applied to the data to measure amplitude and period of the oscillations.

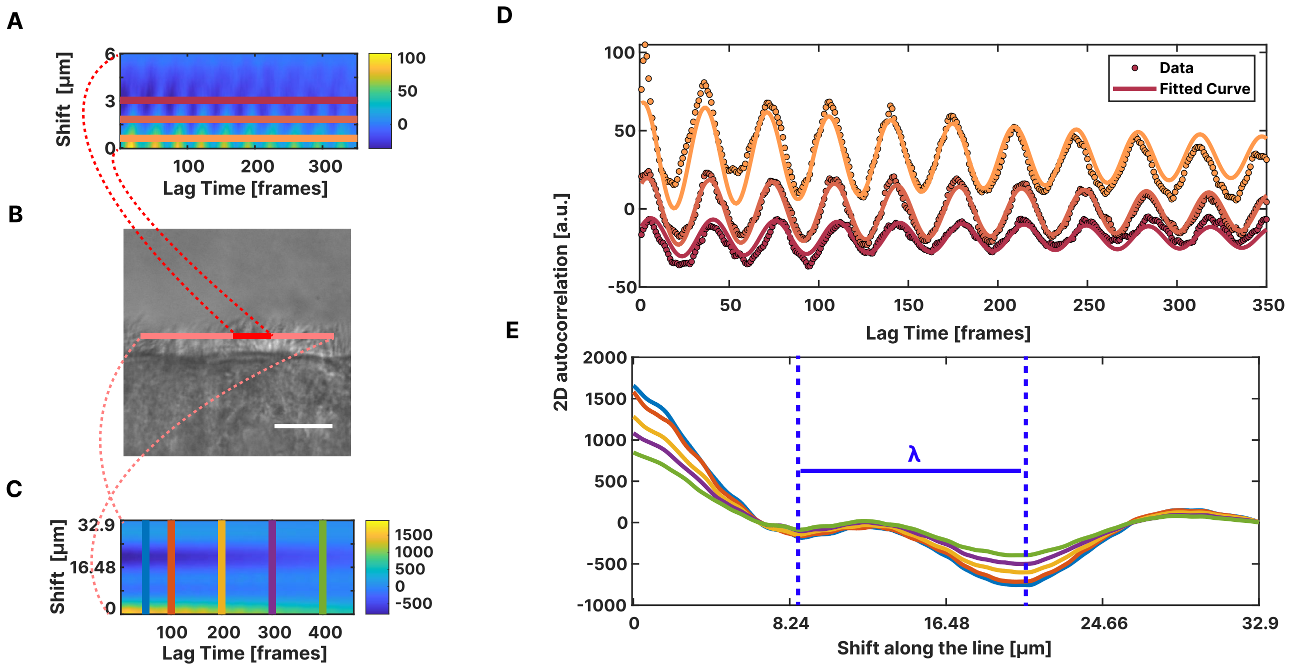


**Fig. S11. MW speed and wavelength measurements.** Grey-scale intensities measured on both short (100 pixels) and long (up to 1000 pixels) segments within the ciliary layer were cross-correlated to determine MW properties and ciliary coordination length-scale, respectively. We examined the autocorrelation in relation to lag time at specific space shifts (A,D). Data points are fitted using the equation of a damped oscillator (equation S5). MW speed is determined by analysing the phase lag between autocorrelations across various space shifts. Further, we extended the segment and we looked at the evolution of the autocorrelation at fixed lag times (C). All the autocorrelation vs space shifts goes to zero and doesn't recover up, meaning we are recording one wavelength in the FOV (E). The wavelength $\lambda$ is measured as the distance between two consecutive peaks (or valleys) in this plot. The frame rate of this recorded video is 150 fps.

fitted according to the model previously described. The fitted curves are notably in phase, displaying a small average phase shift of $\Delta\Phi = 0.37 \pm 0.24$.

**B. Metachronal wave wavelength.** To assess MW wavelength, we extended our analysis to the maximum segment length within the cilia patch. This is shown in pink in figure S11.B where it measures 500 pixels and in red in figure S12.G where it measures 280 pixels. The measurement of the MW wavelength was achieved by examining autocorrelation against space shift at specific lag times (50, 100, 200, 300, and 400 frames). As shown in figure S11.E and figure S12.I, we determined the wavelength $\lambda$ as the distance between two consecutive peaks (or valleys) in the autocorrelation vs space shift plots. The measured wavelength averaged $13.4 \pm 0.8\ \mu m$ across the two experiments.

### 8. Simulations

We compute the flow field around multiple beating ciliary filaments modelled as rigid rods with a length asymmetry. The hydrodynamics of a filamentous structure like a cilium are best described using slender-body theory (SBT) (1, 2). Within this framework, the velocity $\boldsymbol{u}$ of the centreline $C_f$ of a slender filament is related to the hydrodynamic force per unit length, $\boldsymbol{f}_h$, acting on it through the integral equation,

$$\boldsymbol{u}(\boldsymbol{x}_0) = -\frac{1}{8\pi\mu}\boldsymbol{\Lambda}[\boldsymbol{f}_h](\boldsymbol{x}_0) - \frac{1}{8\pi\mu}\boldsymbol{K}[\boldsymbol{f}_h](\boldsymbol{x}_0), \quad \text{[S7]}$$

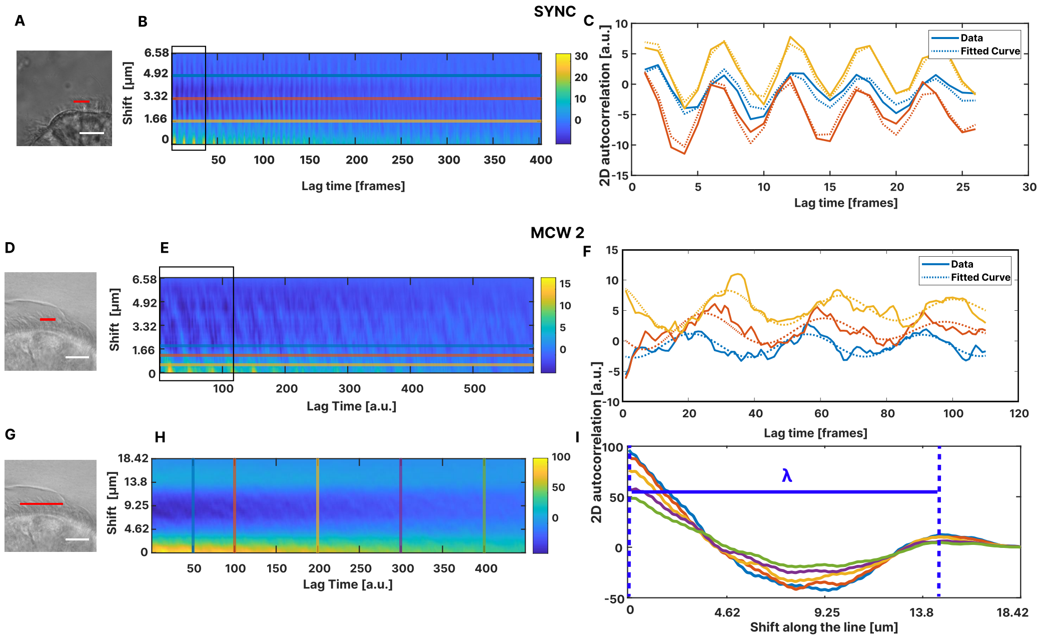


**Fig. S12.** Additional analysis of autocorrelation plots for in-phase beating cilia (A-C) and cilia beating with a constant phase lag - antiplectic methachronal waves - (D-I). The frame rate of the recorded videos are 120 and 150 fps for the synhcronous and MW case, respectively.

where $\boldsymbol{x}_0 = \boldsymbol{x}_0(s,t)$ and $s \in [0, L]$. The no-slip boundary condition on the rigid wall is satisfied by using appropriate image singularities. The local, $\boldsymbol{\Lambda}$, and non-local, $\boldsymbol{K}$, operators are given by

$$\boldsymbol{\Lambda}[\boldsymbol{f}_h](\boldsymbol{x}_0) = [-c(\boldsymbol{I} + \hat{\boldsymbol{s}}\hat{\boldsymbol{s}}) + 2(\boldsymbol{I} - \hat{\boldsymbol{s}}\hat{\boldsymbol{s}})] \cdot \boldsymbol{f}_h(\boldsymbol{x}_0), \qquad \text{[S8]}$$

$$\boldsymbol{K}[\boldsymbol{f}_h](\boldsymbol{x}_0) = \int_0^L \left( \boldsymbol{f}_h(\boldsymbol{x}(s')) \cdot \boldsymbol{G}(\boldsymbol{x}(s'), \boldsymbol{x}_0(s)) - \frac{\boldsymbol{I} + \hat{\boldsymbol{s}}(s)\hat{\boldsymbol{s}}(s)}{|s - s'|} \cdot \boldsymbol{f}_h(\boldsymbol{x}_0(s)) \right) \mathrm{d}s'(\boldsymbol{x}), \qquad \text{[S9]}$$

where $c = \log(\varepsilon^2 e)$. The tensor $\boldsymbol{G}(\boldsymbol{x}, \boldsymbol{x}_0)$ is the free space Green's function for Stokes equation also called the Oseen–Burgers tensor, representing fluid flow produced by a point force,

$$\boldsymbol{G}(\boldsymbol{x}, \boldsymbol{x}_0) = \frac{\boldsymbol{I}}{|\boldsymbol{x} - \boldsymbol{x}_0|} + \frac{(\boldsymbol{x} - \boldsymbol{x}_0)(\boldsymbol{x} - \boldsymbol{x}_0)}{|\boldsymbol{x} - \boldsymbol{x}_0|^3}, \qquad \text{[S10]}$$

where $\boldsymbol{I}$ is the $3 \times 3$ identity tensor. To account for the no slip velocity condition on the wall, we make use of the half-space Green's function, $\boldsymbol{G}^w(\boldsymbol{x}, \boldsymbol{x}_0)$ (3). Let us consider a Stokeslet placed at a distance $h$ above the wall at $y = 0$ such that its location is $(y_1, y_2, h)$. The image singularities are then accordingly located below the wall at $(y_1, y_2, -h)$. The Green's function due to the Stokeslet with its images at an evaluation point $(x_1, x_2, x_3)$ is

$$G^w_{ij} = \frac{\delta_{ij} + \hat{r}_i\hat{r}_j}{r} - \frac{\delta_{ij} + \hat{R}_i\hat{R}_j}{R} + 2h\Delta_{jk}\frac{\partial}{\partial R_k}\left(\frac{h\hat{R}_i}{R^2} - \frac{\delta_{i1} + \hat{R}_i\hat{R}_1}{R}\right), \qquad \text{[S11]}$$

where the vector pointing from the Stokeslet location to the evaluation point is $r_i = (x_1 - y_1, x_2 - y_2, x_3 - h)$, the vector pointing from the image location to the evaluation point is $R_i = (x_1 - y_1, x_2 - y_2, x_3 + h)$ and the matrix $\Delta_{jk}$ is

$$\Delta_{jk} = \begin{bmatrix} 1 & 0 & 0 \\ 0 & 1 & 0 \\ 0 & 0 & -1 \end{bmatrix}. \qquad \text{[S12]}$$

Note that in this slender body theory formulation, the cross-sectional radius of the body varies slowly as $r(s) = 2\varepsilon\sqrt{s(L-s)}$ where $\varepsilon = r(L/2)/L$, ensuring algebraically-accurate results. The cross-sectional radius at the midpoint $s = L/2$ is taken to be equal to the radius of the flagellum, i.e. $r(L/2) = \rho$ . The kernel in Eq. (S9) becomes formally singular when $s = s'$ and this singularity is removed by regularising the integral.

Discretisation the rod into straight elements, and solving the integral equation Eq. (S7), we obtain a linear system of equation $\boldsymbol{u} = \boldsymbol{A}\boldsymbol{f}$, where $\boldsymbol{A}$ is a matrix that contains the Green's function and hence depends on the geometry. Solving this system of equation gives us the desired traction on the rod which we can use to find the flow field anywhere in the half-space.

$$\boldsymbol{u}(\boldsymbol{x}_0) = \int_0^L \boldsymbol{f}_h(\boldsymbol{x}(s')) \cdot \boldsymbol{G}(\boldsymbol{x}(s'), \boldsymbol{x}_0(s))\, \mathrm{d}s'(\boldsymbol{x}), \qquad \text{[S13]}$$

where now $\boldsymbol{x}_0$ can be anywhere in the three-dimensional domain. In the simulations, we create a cubic grid and compute the velocities at each of the grid points.

Each cilium oscillates between back and forth around the mean $\theta = \pi/2$ (perpendicular to the wall). The desired asymmetry in beating is obtained by decreasing the length of the cilium from $L^* = 1$ in the power stroke to $L = 0.6^*$ in the recovery stroke. The velocity at each point of the cilium is then given by rigid body rotation $\Omega$,

$$\boldsymbol{u}(\boldsymbol{x}_0) = \Omega(t)\boldsymbol{e}_y \times (\boldsymbol{x}_0 - \boldsymbol{x}_J) \qquad \text{[S14]}$$

where $\boldsymbol{x}_J$ is the point of attachment of the cilium on the wall. The angle made by the cilia with the base is a smooth function of time as it switches direction when the cilia alternates between recovery and power stroke,

$$\theta_i = 0.5\pi + \frac{1 - \arccos\left(0.99 \sin\left(2\pi(t + \phi_i) + 0.5\pi\right)\right)}{0.5\pi} \cdot \frac{1}{0.9099} \qquad \text{[S15]}$$

where $\phi$ is the phase of an individual cilium, $i$, in the array. The angular velocity is found by taking the derivative of $\theta_i$,

$$\Omega_i = \frac{4.35213 \cos\left(0.5\pi + 2\pi(t + \phi_i)\right)}{\sqrt{1 - 0.9801\left(\sin\left(0.5\pi + 2\pi(t + \phi_i)\right)\right)^2}}. \qquad \text{[S16]}$$

Note that the time at which the simulations are started is arbitrary (corresponding to the initial configuration of the cilia). Hence, the results in the main text are computed by taking the average over one time period from $t \in [0, 1]$. Equations for multiple cilia follow in a straightforward manner. When considering $\boldsymbol{x}_0$ on filament labelled $\alpha$, we simply have nonsingular integrals over remaining $N_{\mathrm{fil}} - 1$ filaments labelled $\beta$.

$$\boldsymbol{u}(\boldsymbol{x}_{0,\alpha}) = -\frac{1}{8\pi\mu}\boldsymbol{\Lambda}[\boldsymbol{f}_h](\boldsymbol{x}_{0,\alpha}) - \frac{1}{8\pi\mu}\boldsymbol{K}[\boldsymbol{f}_h](\boldsymbol{x}_{0,\alpha}) - \sum_{\beta=1,\beta\neq\alpha}^{N_{\mathrm{fil}}} \frac{1}{8\pi\mu}\int_0^L \boldsymbol{f}_h(\boldsymbol{x}(s')) \cdot \boldsymbol{G}(\boldsymbol{x}(s'), \boldsymbol{x}_0(s))\, \mathrm{d}s'(\boldsymbol{x}), \qquad \text{[S17]}$$

The traction (force per unit length) on each filament is a three-dimensional vector. Let each filament be divided into $N_{\mathrm{elm}}$ elements. Hence, for a single filament, we have $3N_{\mathrm{elm}}$ unknowns, and for $N_{\mathrm{fil}}$ we have a total of $3N_{\mathrm{elm}}N_{\mathrm{fil}}$ unknowns. Once these unknowns are determined by solving the integral equation Eq. (S17) numerically, we can determine the flow field anywhere in the domain, by computing an integral similar to equation Eq. (S13) but looping over all the filaments,

$$\boldsymbol{u}(\boldsymbol{x}_0) = \sum_{\alpha=1}^{N_{\mathrm{fil}}} \int_0^L \boldsymbol{f}_h(\boldsymbol{x}(s')) \cdot \boldsymbol{G}(\boldsymbol{x}(s'), \boldsymbol{x}_0(s))\, \mathrm{d}s'(\boldsymbol{x}), \qquad \text{[S18]}$$

where $\boldsymbol{x}_0$ loops over a cubic grid. When computing the trajectories of particles (pathlines), we simply compute the flow velocity at the position of the particle at $\boldsymbol{x}_0$ and advance the particle position in time numerically,

$$\frac{\mathrm{d}\boldsymbol{x}_0}{\mathrm{d}t} = \boldsymbol{u}(\boldsymbol{x}_0). \qquad \text{[S19]}$$

Trajectories of particles released at increments of 0.1 above the epithelial wall and the plane $y = 0.1$ parallel to the beating plane are shown in Fig. S13. The dual slope in the displacement parallel to the epithelium is still observed for particles that get expelled. However, particles close to wall do not get sucked towards the wall as strongly. Hence, their $x-$ trajectories do not exhibit a dual slope. In experiments, the sharp change in slope occurs much earlier which is not seen here.

**Movie S2.** Streamline around an array of 41 cilia with 2 metachronal waves.

1. RE Johnson, An improved slender-body theory for stokes flow. *J. Fluid Mech.* **99**, 411–431 (1980).
2. AK Tornberg, MJ Shelley, Simulating the dynamics and interactions of flexible fibers in stokes flows. *J. Comp. Phys.* **196**, 8–40 (2004).
3. JR Blake, A note on the image system for a stokeslet in a no-slip boundary in *Proc. Camb. Phil. Soc.* (Cambridge University Press), Vol. 70, pp. 303–310 (1971).

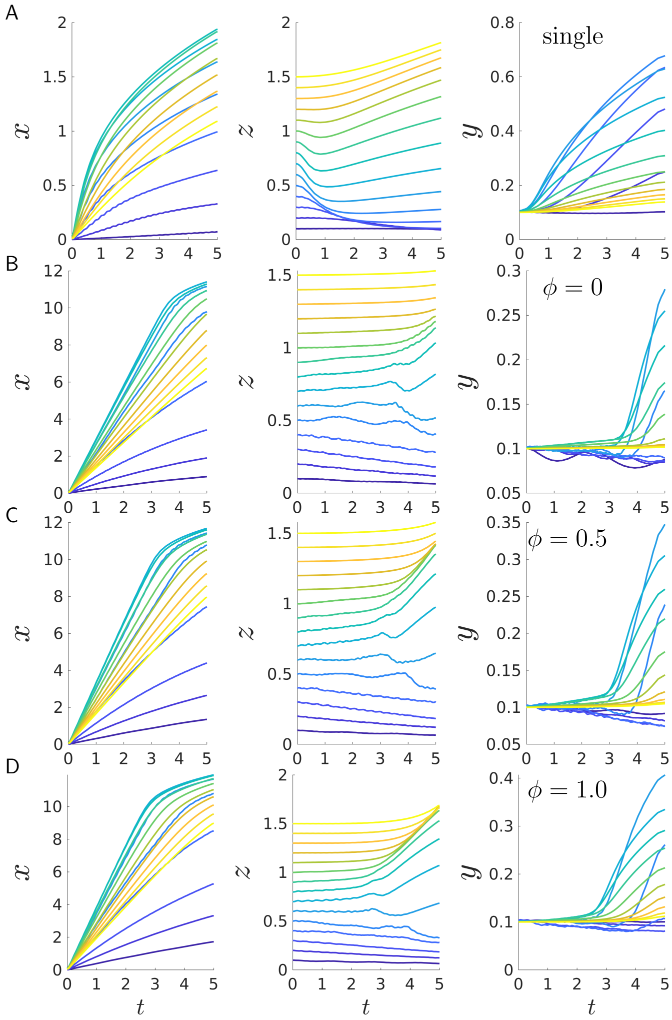


**Fig. S13.** Simulations of particles trajectories released at a height of $z \in [0.1 - 1.5]$ in increments of 0.1 and parallel to $x - z$ plane at $y = 0.1$, in the presence of a single cilium (A), and driven by an array of cilia with varying phase difference, $\phi = 0$ in synchrony (C), $\phi = 0.5$ (D), and $\phi = 1$ (E) and when cilia are beating randomly (F). Particle trajectories in the direction of beating plane in $x-$ and perpendicular to the wall in $z-$ direction are plotted panels (B) and (C). Particle trajectories in the $x - z$ plane are shown in panels (D). In the plot, the spatial simulation unit (s.u.) correspond to the length of an extended cilium in the array. The time interval explored has been chosen to be similar to the one of experiments – cilia beat at $\sim 5$ Hz and the video captured are $\sim 5$ s long so we imaged cilia beating for $\sim 25$ beating cycles.
